# Nascent RNA profiling reveals regulation of gene transcription through productive reiterative initiation in bacteria

**DOI:** 10.1101/2024.12.23.630015

**Authors:** Zhe Sun, Guozhong Huang, Yuqiong Zhang, Shuang Wang, Mikhail Kashlev

**Author notes:** Correspondence: Zhe Sun, Shuang Wang, Mikhail Kashlev.

## Abstract

Productive reiterative initiation, an alternative transcription process accompanying RNA slippage relative to the template DNA and RNA polymerase, results in the incorporation of additional nucleotides into transcripts. However, a comprehensive understanding of the mechanisms underlying productive reiterative initiation and its functional implications has been hindered due to the complexity of heterogenous slippage transcripts. Here, we develop and employ 5’ Terminal Native Elongating Transcript Sequencing (5’TNET-seq) to identify and quantify productive reiterative initiation events in *Escherichia coli*. 5’TNET-seq reveals that over 51% of promoters exhibit productive reiterative initiation. The conserved promoter – 10 region, an appropriate spacer between the –10 region and the TSS, and the transcription initiation region associated with weak RNA-DNA hybrid stability, particularly “AAA” and “TTT” trinucleotide tracts, contribute to high productive reiterative initiation. In addition, up to four nucleotides can be added in a single cycle of productive reiterative initiation. A smaller transcription bubble is observed during productive reiterative initiation, which may stabilize the transcription initiation complex to regulate gene transcription. Our results suggest that productive reiterative initiation emerges as an inherent transcription process regulating biological processes, including cell wall synthesis, independent of protein regulators.

## INTRODUCTION

Reiterative transcription, also known as transcript slippage, represents a noncanonical transcription process generating heterogeneous transcripts through the addition of extra nucleotides^1,2^. In the irregular process, RNA polymerase (RNAP) does not translocate relative to the template DNA. Instead, the nascent RNA slips backward relative to the template DNA, allowing RNAP to repetitively add nucleotides to the 3’ end of the nascent RNA^3–5^. Despite the expectation that regularly transcribed RNA should accurately base-pair with the template DNA strand, RNA-DNA mismatches occur at a substantial rate, exhibiting sequence preferences^6–8^. Reiterative transcription, as an alternative transcription process, adds extra nucleotides to the RNA 3’ end, which may result in mismatches when aligned with the genome. This process is prevalent across various species from prokaryotes to eukaryotes, including RNA and DNA viruses^9–12^, *Escherichia coli*^13^, *Bacillus subtilis*^14^, *Saccharomyces cerevisia*e^15^ and humans^16,17^.

Amino acid mutations in the RNAP β subunit and the bacteriophage λN protein have been reported to influence the slippage rate by affecting the stability of the RNA-DNA hybrid^18–20^. The low stability of the RNA-DNA hybrid in the ternary transcription complex (DNA-RNA-RNAP) leads to the extensive occurrence of reiterative transcription in distinct transcription phases, including transcription initiation, elongation, and termination. Notably, reiterative transcription during elongation has been extensively investigated, driven by mutations introduced at slippery sites predominantly within open reading frames (ORFs)^21^. The resultant frameshift protein mutants generated by reiterative transcription may undergo truncation, leading to loss of function, or conversely, restoration to functional proteins, such as the membrane protein TssM and transposase^22,23^. Subsequent to the elongation phase, reiterative transcription induced by RNA slippage can also stimulate transcription termination, manifested by misalignment between the RNA and DNA templates in the poly(U) tract proximal to the stem-loop structures of intrinsic terminators^24,25^.

Furthermore, reiterative transcription has been observed during the initiation phase (termed as “reiterative transcription initiation”) at several promoters of the *pyrBI*, *codBA* and *upp* genes^2,13,26^. This initiation process regulates gene expression in response to intracellular signals. In this process, RNA engages in a short 2–3-bp hybridization with the template DNA at a homopolymeric tract immediately downstream of the transcription start site (TSS) and subsequently slips relative to the template^27^. The nascent RNA undergoes repetitive cycles of elongation by RNAP, adding the same nucleotide, which may extend toward σR3.2 of the housekeeping σ factor σ^70^ as in regular transcription initiation^28^. This process is triggered directly by elevated UTP concentration at the *E. coli pyrBI* promoter or indirectly through TSS switching stimulated by excessive UTP at the *codBA* and *upp* promoters^2,13,26^, leading to the addition of a homopolymeric tract to the RNA and the subsequent release of nascent RNA^3,4^, referred to herein as “non-productive reiterative initiation”.

Despite the detachment of slippage RNA transcripts from the transcription initiation complex, similar with abortive transcripts^29^, RNA transcripts from the *pyrG* promoter of *B. subtilis* exiting through the RNAP main channel^28,30^ could switch to the RNA-exit channel before dissociation. These transcripts can revert to regular transcription initiation during this process, in contrast to the *E. coli pyrBI*, *codBA*, and *upp* promoters, a phenomenon termed “productive reiterative initiation”. The occurrence of productive reiterative initiation in vivo remains largely unknown due to limited investigation and a lack of in vivo evidence. High-throughput sequencing methods, specifically dRNA-seq and Send-seq, capable of determining transcription start sites^31,32^, have indicated the presence of RNA-DNA mismatches near the TSSs of some promoters. Additionally, the 5’ end of many promoter-proximal reads from RNET-seq, designed to identify transcription pause sites (PSs), exhibited mismatched nucleotides^33^. A reasonable assumption arises that these mismatches, derived from indels, are caused by productive reiterative initiation rather than conventional transcription errors. Despite the examination of transcript slippage at the *lacCONS* promoter using massively systematic transcript end readout (MASTER)^34^, the universality, mechanism, and biological significance of productive reiterative initiation in bacteria remain substantially unexplored.

In this study, we developed a novel method termed 5’ Terminal Native Elongating Transcript Sequencing (5’TNET-seq) for the comprehensive identification and quantification of genome-wide productive reiterative initiation in vivo within *E. coli*. The application of 5’TNET-seq unveiled a prevailing occurrence of productive reiterative initiation at numerous promoters, many of which exhibited a notable high slippage ratio, estimated as a ratio of productive reiterative initiation transcripts to the total transcripts originating from a promoter. Subsequent analysis revealed the involvement of the promoter –10 region, the spacer between the –10 region and the TSS, and the transcription initiation region in facilitating productive reiterative initiation. Particularly, the presence of a “T” or “A” tract in the transcription initiation region significantly elevates the cycle number of repetitive nucleotide addition. Strikingly, our findings reveal that productive reiterative initiation enhances the stability of the transcription initiation complex, promotes promoter escape by inducing a clash between σR3.2 and the nascent RNA 5’ end, thereby augmenting gene transcription and regulating cell wall synthesis.

## RESULTS

### Identification of obscure TSSs and transcription pause sites through 5’TNET-seq

To comprehensively unravel genome-wide productive reiterative initiation in *E. coli*, we developed the **5’ t**erminal **n**ative **e**longating **t**ranscript **seq**uencing (5’TNET-seq). In essence, this technique involves the extraction and size-fractionation of nascent RNA from elongating ternary complexes. Subsequently, short nascent RNAs (≤ 100-nt) are purified, treated with 5’-monophosphate-dependent terminator exonuclease, and utilized for the construction of high-throughput sequencing libraries concurrently (Fig. 1a). Compared to dRNA-seq, which utilizes total RNA as the starting material for library construction (Supplementary Fig. 1), 5’TNET-seq offers superior resolution for detecting productive reiterative initiation. Conversely, RNET-seq, which identifies pause sites, may eliminate the products of productive reiterative initiation due to potential digestion of the nascent RNA 5’ end by RNase I in promoter-proximal regions (Supplementary Fig. 1). The resulting 5’TNET-seq profiles consistently identified TSSs and pause sites simultaneously, as validated by dRNA-seq and RNET-seq, respectively (Fig. 1b). Comparative analysis demonstrated that approximately 70% of promoter-proximal pauses identified by RNET-seq were also captured by 5’TNET-seq, exhibiting a similar RNAP pausing pattern (Fig. 1c, Supplementary Fig. 2), thus affirming the efficacy of 5’TNET-seq in PSs detection.

**Fig. 1.**
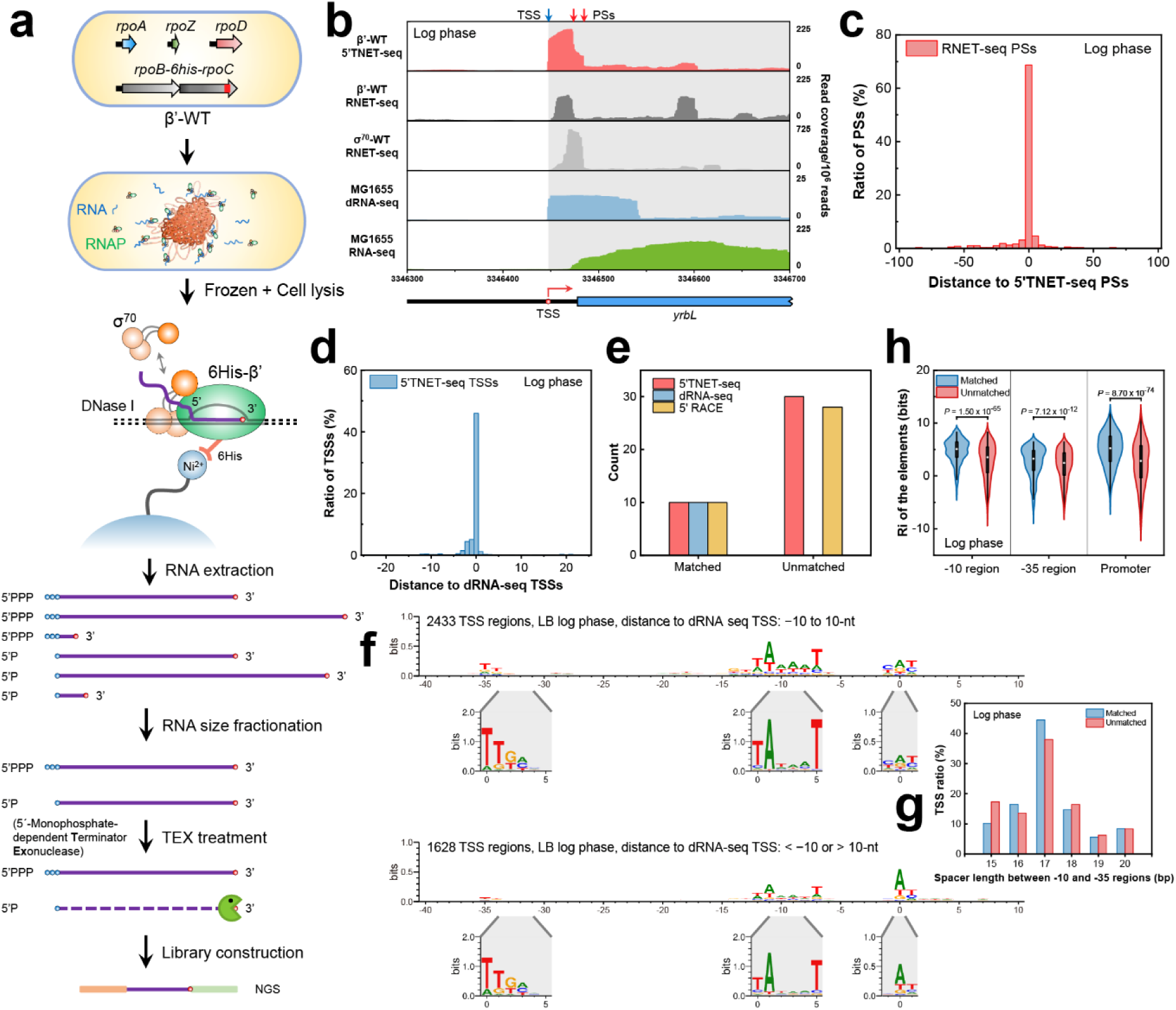
Development of 5’TNET-seq for identifying obscure TSSs and transcription pause sites. **a** Schematic representation of the 5’TNET-seq workflow. This technique involves isolating native elongating ternary complexes, extracting nascent RNA, size fractionation, and 5’-monophosphate-dependent terminator exonuclease processing. The resultant RNA is utilized for library construction and high-throughput sequencing. **b** Comparative analysis of 5’TNET-seq, RNET-seq, dRNA-seq, and RNA-seq profiles near the *yrbL* TSS during *E. coli* log phase. **c** Examination of pause sites identified by 5’TNET-seq compared to those identified by RNET-seq. **d** Evaluation of TSSs detected by 5’TNET-seq in contrast to those identified by dRNA-seq. TSSs identified by 5’TNET-seq that are within a 50-bp region centered on the nearest TSSs detected by dRNA-seq are shown. **e** 5’ RACE validation of TSSs discovered by 5’TNET-seq, categorized by distances, either ≤ 10-nt (matched, n = 2433) or > 10-nt (unmatched, n = 1628) relative to the TSSs identified by dRNA-seq. **f** Sequence logos illustrating the regions surrounding “matched” (top) and “unmatched” (bottom) TSSs discovered by 5’TNET-seq. Corresponding sequence logos depicting the aligned –35, –10, and TSS regions are displayed below. **g** Comparative analysis of spacer lengths between the –10 and –35 regions. **h** Violin plot assessment of information content in the –10 region, –35 region, and promoter sequences for “matched” (n = 2433) and “unmatched” (n = 1628) TSSs. In this and all subsequent violin plots, the box represents the interquartile range, the whiskers extend to 1.5 times the interquartile range, and the white cycle indicates the median. Statistical analysis was executed employing a two-tailed Mann-Whitney *U*-test.

In comparison with dRNA-seq, it was observed that only ∼46% of 5’TNET-seq TSSs matched those identified by dRNA-seq, which may be partially attributed to productive reiterative initiation (Fig. 1d). Subsequent analysis categorized 5’TNET-seq TSSs into two groups: “matched” and “unmatched”, based on whether the distances from the nearest TSS identified by dRNA-seq are ≤ 10-nt or > 10-nt, respectively. The existence of “unmatched” TSSs was unequivocally confirmed through 5’ rapid amplification of cDNA ends (5’ RACE), underscoring the precision of 5’TNET-seq (Fig. 1e, Supplementary Fig. 3). Sequence logos indicated a consistent spacer length between the –10 region and the –35 region (Fig. 1f and g), while revealing lower conservation of the –10 region and –35 region in “unmatched” promoters (Fig. 1h), supported by information content quantified by the σ^70^ promoter model^35^. This observation was consistent during the stationary phase (Supplementary Fig. 4), highlighting 5’TNET-seq’s capability in unveiling less conserved promoters. This capability proves beneficial for the comprehensive and precise analysis of productive reiterative initiation.

### Genome-wide identification and quantification of productive reiterative initiation

In the pursuit of comprehensively understanding of productive reiterative initiation, 5’TNET-seq reads with 5’ end mismatches due to additional nucleotide synthesis at the 5’ end of nascent RNA during productive reiterative initiation were investigated. We extracted all reads with a 5’ end mismatch from 5’TNET-seq reads that uniquely mapped to the genome. Approximately 7% of these uniquely mapped reads exhibited 5’ end mismatches, notably enriched with “TT” (Fig. 2a, top). We then designated the nucleotide immediately downstream of the 5’ end mismatches as the transcript origin and compared it with dRNA-seq TSSs. Intriguingly, around 19% of TSSs identified by dRNA-seq displayed nascent RNAs with 5’ end mismatched nucleotides (Fig. 2a, bottom). The counts of 5’TNET-seq reads originating from dRNA-seq TSSs with 5’ end mismatches were significantly higher than those from randomly selected sites (Fig. 2b), suggesting that these mismatches were not artifacts introduced during library construction or sequencing, implying a potential genome-wide occurrence of productive reiterative initiation.

**Fig. 2.**
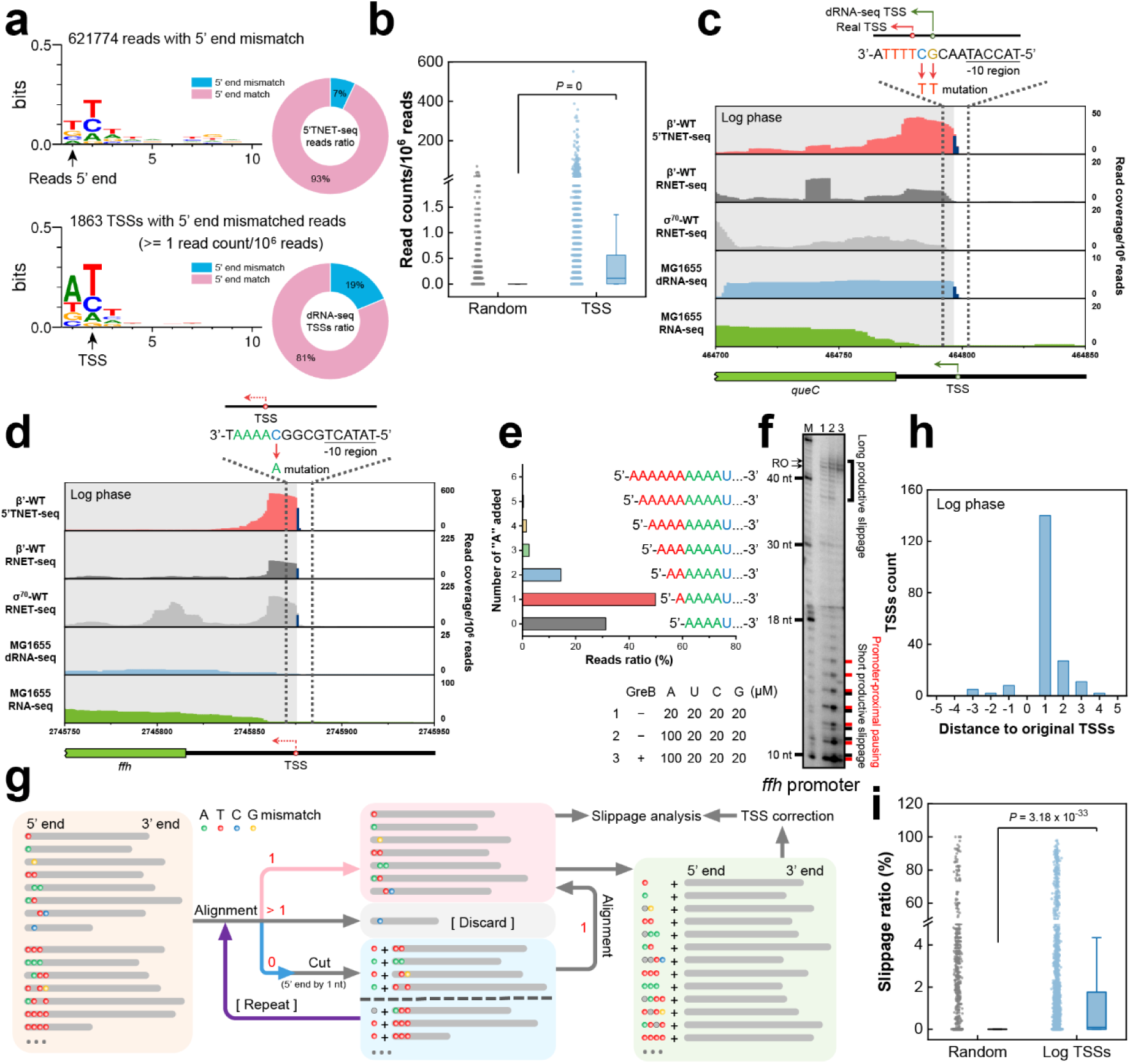
Identification and quantification of productive reiterative initiation in vivo using 5’TNET-seq. **a** Sequence logos depict the 5’TNET-seq reads characterized by 5’ end mismatches and the transcription initiation regions identified in dRNA-seq with 5’ end mismatched reads. **b** Boxplot presenting the counts of 5’ end mismatched reads originating from regions within 5-nt of the dRNA-seq TSSs in comparison to randomly selected sites (n = 9966). In this and all subsequent boxplots, the box corresponds to the interquartile range, the line denotes the median, and the whiskers extend to 1.5 times the interquartile range. A two-tailed Mann-Whitney *U*-test was conducted for statistical analysis. Exemplified 5’TNET-seq profiles pertaining to the *queC* **(c)** and *ffh* **(d)** TSS regions, characterized by notably elevated ratios of 5’ end mismatched reads. The blue bars within these profiles represent the nucleotides displaying 5’ end mismatches in the sequencing reads. **e** Overview of 5’TNET-seq reads exhibiting a variable number of “A” nucleotides added to the 5’ end of the *ffh* transcripts. **f** In vitro transcription assays validating productive reiterative initiation at the *ffh* promoter. Specific conditions for each lane are detailed on the left side of the panel. M, marker; RO, run-off product. **g** The processing pipeline for 5’TNET-seq reads, proficient in extracting all reads stemming from productive reiterative initiation and calculating the slippage ratios and patterns. **h** Distribution of corrected TSSs relative to the original TSSs following pipeline analysis. **i** Boxplot illustrating the comparative analysis of the slippage ratios across all TSSs and an equivalent number of random sites (n = 4188), with statistical analysis performed using a two-tailed Mann-Whitney *U*-test.

A representative 5’TNET-seq profile at the *queC* promoter, characterized by a T-track, exhibited a high 5’ end mismatch ratio, consistent with observations from dRNA-seq (Fig. 2c). This pattern extended to other promoters, such as *gcvT* and *greA*, containing A-tracks, with 5’ end mismatched nucleotides confirmed by 5’ RACE (Supplementary Fig. 5). However, the addition of 5’ end nucleotides during productive reiterative initiation posed challenges to algorithms predicting TSSs, leading to inaccuracies (Fig. 2c, Supplementary Fig. 5). Notably, newly identified TSSs by 5’TNET-seq, like the TSS at the *ffh* promoter, exhibited a high ratio of mismatched nucleotides right upstream of the TSS (Fig. 2d). Because sequencing reads with more than two mismatches were unable to be aligned to the genome, we manually counted the number of additional “A” nucleotides added to the *ffh* nascent RNA. It’s intriguing that reads with one “A” added outnumbered perfectly aligned reads, and up to six “A” nucleotides could be added to the *ffh* nascent RNA (Fig. 2e). To further verify the existence of productive reiterative initiation, in vitro transcription assay with the *ffh* promoter was performed and identified short and long productive slippage products with increased slippage products at higher ATP concentration, which were eliminated by GreB addition (Fig. 2f). This underscores the occurrence of productive reiterative initiation during transcription initiation and the possible role of the transcription elongation factor GreB in suppressing it.

To delve into the mechanism of this widespread productive reiterative initiation, we devised a processing pipeline to correct ambiguous 5’TNET-seq reads with 5’ end nucleotide additions (Fig. 2g). Basically, reads uniquely aligned to the genome were retained, while unaligned reads underwent iterative 1-nt trimming from the 5’ end until they were either uniquely aligned or discarded due to multiple alignments. The original 5’ end of the nascent RNA before productive reiterative initiation was identified, and the trimmed 5’ nucleotides were collected to quantify productive reiterative initiation. Post-processing, the coordinates of approximately 200 TSSs were corrected, mostly shifting downstream by 1–2-nt during both the log and stationary phases due to productive reiterative initiation (Fig. 2h, Supplementary Fig. 6a). The slippage ratio (mismatched reads/total aligned reads × 100%) calculated for each TSS was significantly higher than those for random sites, affirming that productive reiterative initiation is inherent during transcription initiation as opposed to an artifact introduced by sequencing errors (Fig. 2i, Supplementary Fig. 6b, Supplementary Table 1). Further analysis showed that 51.6% and 72.6% of promoters identified by 5’TNET-seq showed productive reiterative initiation in the log and stationary phases, respectively (Supplementary Fig. 7).

### Identification of promoter elements influencing productive reiterative initiation

During the log phase, we systematically categorized TSSs into four distinct groups based on their corresponding slippage ratios (SR): “high” (40%–100%), “medium” (10%–40%), “low” (1%–10%) and “no” (0%). Aligning the DNA regions corresponding to these categories revealed three conserved regions that distinguish each group: the transcription initiation region (TIR, the first three nucleotides from the TSS), the –10 region, and the spacer (nucleotides between the –10 region and the TSS) (Fig. 3a). Sequence analysis of “high slippage” and “medium slippage” TIRs demonstrated that “AAA” and “TTT” trinucleotides were the most prevalent (Fig. 3b), consistent with the sequence logo. Significantly, slippage ratios of “AAA” and “TTT” TIRs were higher than those for other “T” and “A” combinations (–G/C) in TIRs (Fig. 3c). Nevertheless, TIRs lacking “G” or “C” (–G/C) still exhibited higher slippage ratio than those without “A” or “T” (–A/T), indicating a preference for TIRs containing “A” or “T”, particularly “AAA” and “TTT” trinucleotides. In vitro transcription assays validated these finding, showing that promoters featuring “AAA” or “TTT” TIRs had markedly higher slippage ratios compared to other TIRs (Fig. 3d, Supplementary Fig. 8).

**Fig. 3.**
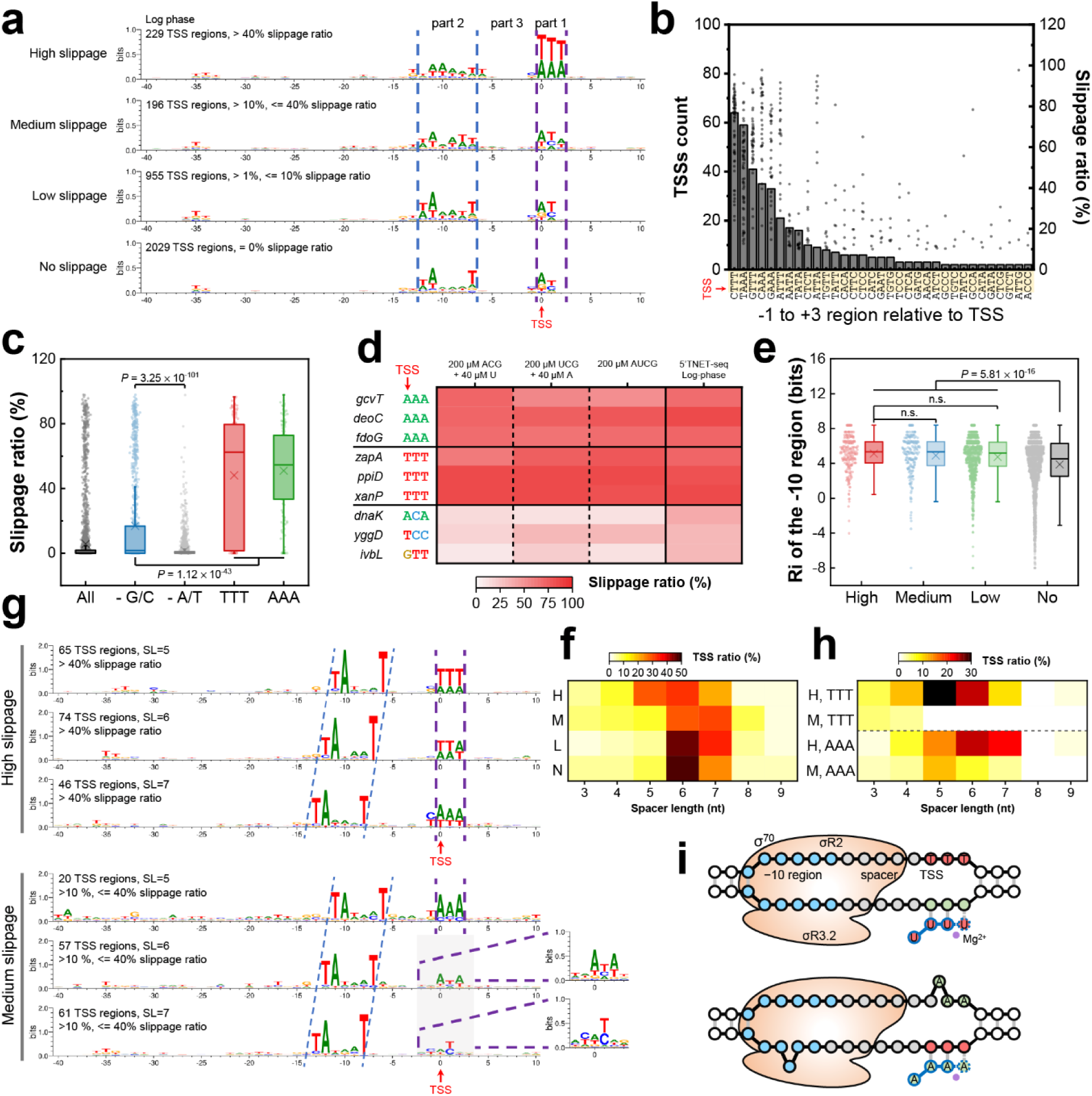
Impact of promoter elements on productive reiterative initiation. **a** Sequence logos illustrating DNA regions encompassing TSSs categorized by slippage ratio. Part 1, transcription initiation region; Part 2, –10 region; Part 3, spacer between the –10 region and the TSS. **b** Histogram presenting the counts and slippage ratio of transcription initiation regions exhibiting productive reiterative initiation. Each dot represents a TSS with high or medium slippage ratios. Areas shaded in yellow denote transcription initiation regions. **c** Boxplot comparing slippage ratios among different patterns of transcription initiation regions. All, all transcription initiation regions (n = 4188); –G/C, regions without “G” and “C” (n = 1138); – A/T, regions without “A” and “T” (n = 3050); TTT, regions with “TTT” (n = 187); AAA, regions with “AAA” (n = 142). Statistical analysis was conducted via a two-tailed Mann-Whitney *U*-test. **d** Heatmap comparing slippage ratios of transcription initiation regions with different patterns through in vitro transcription. NTP concentrations for in vitro transcription assays are indicated at the top. **e** Boxplot comparing information content of the –10 region for TSSs with varying slippage ratios, categorized as high (n = 229), medium (n = 196), low (n = 955) and no (n = 2029) slippage. Statistical analysis was conducted using a two-tailed Mann-Whitney *U*-test. **f** Distribution of spacer lengths in promoters categorized by slippage ratio. H, high slippage; M, medium slippage; L, low slippage; N, no slippage. **g** Sequence logos of promoter regions with high and medium slippage featuring spacer lengths of 5, 6 or 7-nt. **h** Spacer length distribution within promoters featuring “TTT” or “AAA” transcription initiation regions. The promoters demonstrating high (H) and medium (M) slippage are displayed individually. **i** Model of pretranslocated states within promoters featuring “TTT” or “AAA” transcription initiation regions after one single cycle of reiterative transcription. The bulged nucleotide signifies scrunching within the unwound transcription bubble. Blue circles, –10 region; grey circles, spacer between the –10 region and the TSS; pink circles, transcription initiation regions; purple circle, Mg^2+^.

Beyond the TIR, the information content of the –10 region from “high slippage”, “medium slippage”, and “low slippage” promoters was significantly higher than that of the “no slippage” category. This underscores the necessity of a relatively conserved –10 region for fundamental productive reiterative initiation (Fig. 3e). Notably, “high slippage” promoters exhibited a seemingly shifted –10 region, prompting a detailed analysis of their spacer length. It was revealed that “high slippage” promoters favored a 5–6-nt spacer, as opposed to the 6–7-nt spacer observed in “medium slippage” promoters (Fig. 3f, Supplementary Fig. 9). Further classification of “high slippage” and “medium slippage” promoters demonstrated that “TTT” TIRs corresponded to a 5-nt spacer, while “AAA” TIRs with a 5-nt spacer fell into the “medium slippage” group (Fig. 3g). A substantial proportion of “high slippage” promoters with “TTT” or “AAA” TIRs exhibited 5-nt or 6–7-nt spacers, respectively (Fig. 3h), indicating that “TTT” or “AAA” TIRs necessitate an adapted spacer length for the generation of high levels of productive reiterative initiation. Although both “TTT” and “AAA” TIRs contributed to most “high slippage” promoters, “TTT” TIRs displayed a significantly shorter spacer length and higher slippage ratio compared to “AAA” TIRs (Supplementary Fig. 10). This leads to no scrunching or 1-bp scrunching within the unwound transcription bubble for promoters with “TTT” or “AAA” TIRs after one cycle of productive reiterative initiation (Fig. 3i).

### Contribution of promoter elements to slippage pattern and cycles of productive reiterative initiation

The nascent RNA originating from the *B. subtilis pyrG* promoter, featuring a transcription initiation region of “GGG”, exhibits consistent 1-nt slippage relative to RNAP and template DNA in each cycle of reiterative transcription initiation. However, our 5’TNET-seq data revealed diverse slippage patterns at “high slippage” and “medium slippage” promoters, encompassing one, two, three, and four nucleotides slippage per cycle of productive reiterative initiation (Fig. 4a). These promoters or TSSs were defined as 1nc, 2nc, 3nc, and 4nc (nc, **n**ucleotides slippage per **c**ycle), respectively. Notably, the number of 2nc, 3nc, and 4nc promoters and their slippage ratios were substantially lower than those of 1nc promoters. An additional feature of reiterative transcription involves the variable addition of extra nucleotides to the RNA 5’ end. Our calculations demonstrated that certain promoters adding one nucleotide per slippage cycle could accumulate over 15 nucleotides, particularly adding more “T” nucleotides to the RNA 5’ end originating from promoters commencing with “TT” transcription initiation regions and having shorter spacers compared to those with “AA” (Fig. 4b, Supplementary Fig. 11a). Further analysis revealed a median addition of 2.0 “T” nucleotides in contrast to 1.2 “A” nucleotides (Fig. 4c). In vitro transcription assays validated the 5’TNET-seq results, affirming increased nucleotide additions at promoters with a “TTT” TIR (Supplementary Fig. 11b and c).

**Fig. 4.**
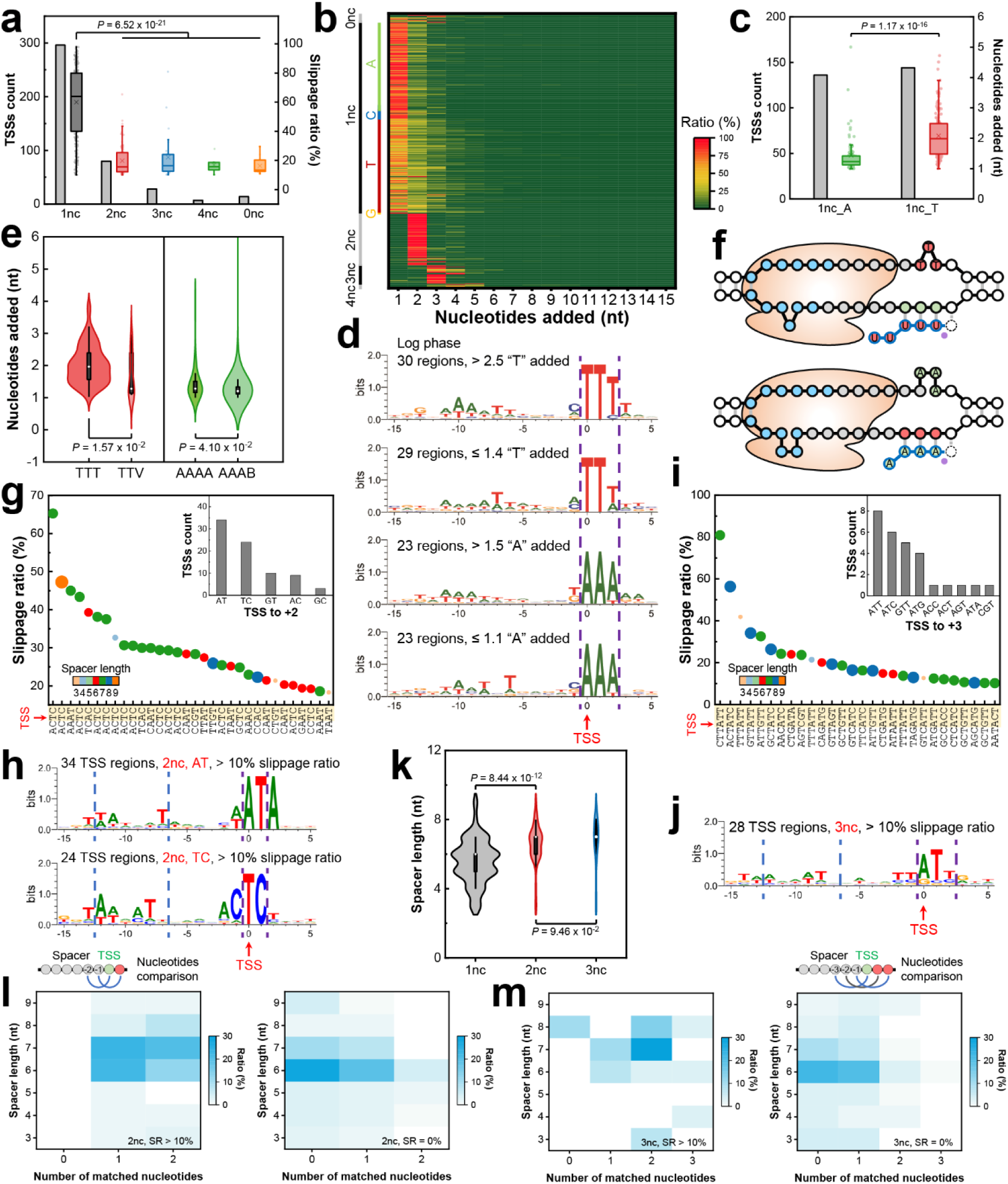
Influence of promoter elements on patterns and cycles of productive reiterative initiation. **a** Boxplot showing the comparative analysis of slippage ratios and counts among promoters across productive reiterative initiation patterns. 1nc, n = 296; 2nc, n = 80; 3nc, n = 28; 4nc, n = 7; 0nc, n = 14. Statistical analysis for the high and medium slippage promoters was conducted via a two-tailed Mann-Whitney *U*-test. **b** Ratio of 5’TNET-seq reads displaying a varying number of nucleotides added to the 5’ end of nascent RNA. Each row corresponds to a TSS. 1nc was further categorized into four groups: AA, CC, TT or GG homopolymeric tract originating from TSS. **c** Boxplot comparing the average number of nucleotides added for the 1nc transcription initiation regions starting with AA or TT. 1nc_A, AA (n = 136); 1nc_T, TT (n = 144). Statistical analysis was performed by a two-tailed Mann-Whitney *U*-test. **d** Sequence logos representing promoter regions with a varying number of “T” or “A” additions. **e** Violin plot comparing the number of nucleotides added for transcription initiation regions with different lengths of “T” or “A” homopolymeric repeats. TTT, n = 126; TTV, n = 18; AAAA, n = 42; AAAB, n = 87. V strands A, C or G; B for C, G or T. A two-tailed Mann-Whitney *U*-test was conducted for statistical analysis to generate the *P* values. **f** Representative states of promoters featuring “TTT” or “AAA” transcription initiation regions after productive reiterative initiation and re-entry into regular transcription initiation. **g, i** Bubble maps illustrating the slippage ratio and spacer length for the 2nc **(g)** and 3nc **(i)** promoters. The 2nc and 3nc transcription initiation regions were ranked by the slippage ratio. The color and size of the bubble indicate the spacer length between the –10 region and the TSS. Only the 2nc promoter region with a slippage ratio exceeding 18% is displayed. Insets show the number of transcription initiation regions originating with the specific dinucleotide or trinucleotide. **h, j** Sequence logos of the 2nc promoter regions that initiate transcription from AT or TC **(h)** and 3nc promoter regions **(j)**. **k** Violin plot comparing the spacer length between 1nc and 2nc, 3nc promoter regions. The statistical analysis utilized a two-tailed Mann-Whitney *U*-test. **l, m** Heatmaps revealing the correlation between the number of matched nucleotides and spacer length for the 2nc **(l)** and 3nc **(m)** promoter regions. The regions with high and medium slippage (left) and those with no slippage (right) were analyzed separately.

To investigate the factors influencing the variable number of nucleotides added at 1nc promoters, we aligned promoters with more or fewer nucleotides added by productive reiterative initiation. It became evident that longer homopolymeric “T” or “A” repeats stimulated more cycles of productive reiterative initiation (Fig. 4d and e, Supplementary Fig. 12a and b). However, the Ri of the –10 region and spacer length did not exhibit significant differences (Supplementary Fig. 12c and d). These findings indicate that the primary determinant for the number of added nucleotides is the length of the homopolymeric repeats in transcription initiation regions. Representative states following re-entry into regular transcription initiation involve the addition of two “U” and one “A”, coupled with 1-bp and 2-bp scrunching for 1nc promoters with “TTT” and “AAA” TIRs (Fig. 4f).

Numerous 2nc and 3nc promoters also exhibited elevated slippage ratios. To elucidate the factors prompting two or more nucleotides slippage in one cycle of productive reiterative initiation, we scrutinized the correlation between slippage ratio and the sequences of transcription initiation regions and spacer length for 2nc and 3nc promoters. Most 2nc promoters has a 7-nt spacer, with “AT” and “TC” TIRs being the most prevalent (Fig. 4g). Sequence logos further highlighted the inclination of these promoters toward a 7-nt spacer. Additionally, nucleotides immediately upstream of the transcription initiation region exhibited a similar pattern to that of the downstream transcription initiation region (Fig. 4h). A similar trend was observed for 3nc promoters, except that the majority of 3nc promoters tended to have an even longer spacer (7–8-nt) (Fig. 4i and j). Statistical analysis confirmed longer spacers in 2nc and 3nc promoters compared to 1nc promoters (Fig. 4k), suggesting the pivotal role of a larger spacer in facilitating more than one nucleotide slippage during productive reiterative initiation. Moreover, a comparison of the two and three nucleotides immediately upstream of the 2nc and 3nc TSSs with the corresponding nucleotides in the transcription initiation regions revealed a higher ratio of matched nucleotides compared to “no slippage” promoters (Fig. 4l and m). These analyses suggest that TIR-simulative DNA sequences immediately upstream of the TSS, coupled with a longer spacer, may contribute to stabilizing multiple nucleotides slippage.

### Regulation of promoter-proximal pausing and gene transcription by productive reiterative initiation

The phenomenon of reiterative transcription has been extensively investigated, primarily focusing on its role in releasing nascent transcripts through the iterative addition of extra nucleotides, ultimately leading to the attenuation of transcription for genes such as *pyrBI*^4^. However, the functional implications of productive reiterative initiation, involving the addition of extra nucleotides to the 5’ end of nascent RNA upon re-entry into regular transcription initiation, remain largely unexplored. Given that 5’TNET-seq enables the simultaneous detection of transcription initiation events and pausing, we evaluated the correlation between productive reiterative initiation and promoter-proximal pausing. Remarkably, our analysis revealed an intriguing relationship between improved slippage ratio and heightened pausing signals in the promoter-proximal region, specifically at 10–30-nt distance downstream of TSS (Fig. 5a). Conversely, promoters featuring identified pause sites in the promoter-proximal regions exhibited significantly higher slippage ratios compared to their counterparts (Fig. 5b). To experimentally validate this correlation, we selected the *zapA* promoter with a “TTT” TIR, characterized by varying numbers of “U” additions according to 5’TNET-seq (Fig. 5c), for in vitro transcription assays. The results confirmed robust productive reiterative initiation (Fig. 5d), consistent with the 5’TNET-seq data. However, the ratio of promoter-proximal pausing products to long productive slippage products exhibited only marginal changes under varying NTP concentrations for both *zapA* and *ffh* promoters tested (Fig. 5e). These results suggest that productive reiterative initiation does not contribute significantly to promoter-proximal pausing. Instead, the pausing is influenced by increased transcription initiation, evident in elevated run-off products with higher UTP concentration and reduced run-off products after mutating the transcription initiation region from “TTT” to “TAT” (Fig. 5d). This observation prompted a hypothesis that productive reiterative initiation might stimulate transcription initiation.

**Fig. 5.**
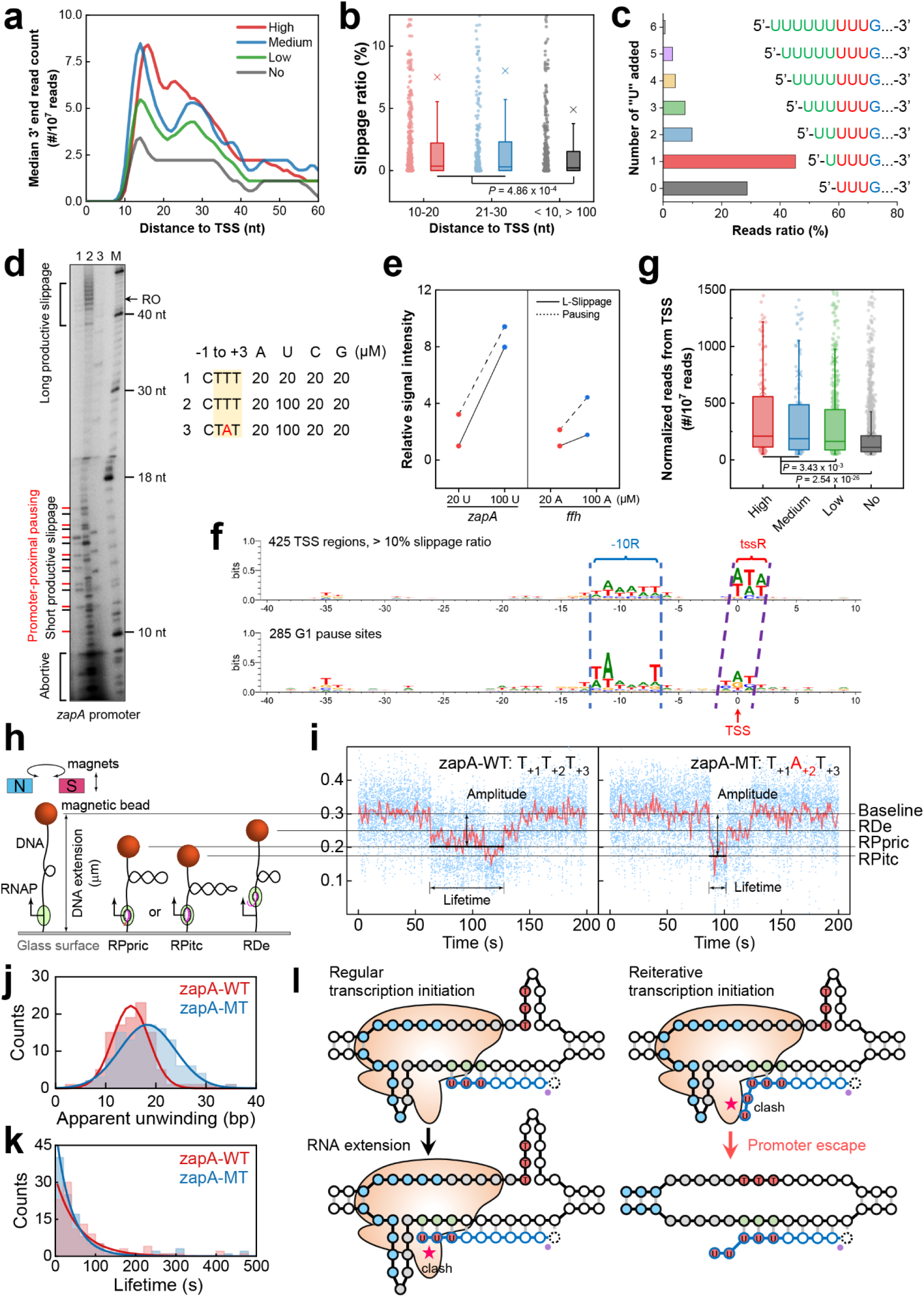
Regulation of productive reiterative initiation on promoter-proximal pausing and gene transcription. **a** Profiles of 3’ end read counts from 5’TNET-seq for promoters with high, medium, low, and no slippage. Median 3’ end read counts were calculated for each coordinate relative to the TSSs in each group. **b** Boxplot comparing the slippage ratio of promoters with transcription pausing in the 10–20-nt (n = 1003) and 21–30-nt (n = 449) regions, or those without transcription pausing in the 11–100-nt (n = 662) regions relative to the TSSs. Statistical differences were assessed using the two-tailed Mann-Whitney *U*-test. **c** Distribution of 5’TNET-seq reads with varying numbers of additional “U” added to the 5’ end of *zapA* transcripts. **d** In vitro transcription assay illustrating productive reiterative initiation and promoter-proximal pausing at the *zapA* promoter. Conditions for each lane are detailed on the right side of the panel. M, marker; RO, run-off product. **e** Relative signal changes of promoter-proximal pausing and long productive slippage products in low and high slippage conditions. The signal intensity of long productive slippage products in low slippage conditions was normalized to 1. **f** Comparison of sequence logos between promoters with high and medium slippage and those with promoter-proximal pausing. **g** Boxplot depicting the comparison of 5’TNET-seq read counts originating from TSSs with high (n = 229), medium (n = 196), low (n = 955) and no (n = 2029) slippage. *P* values were calculated using the two-tailed Mann-Whitney *U*-test. **h** Schematics of single-molecule magnetic trapping assay for transcription experiments under positively supercoiled DNA. RPpric, RNAP-promoter productive reiterative initiation complex; RPitc, RNAP-promoter initial transcribing complex; RDe, RNAP-DNA elongation complex. **i** Typical trajectories representing different transcription bubble sizes of the initiation states: RPpric for zapA-WT construct (left) and RPitc for zapA-MT construct (right), respectively. zapA-WT, wildtype *zapA* promoter; zapA-MT, *zapA* promoter derivate with a single mutation from T_+2_ to A_+2_. Unwinding levels (**j**) and lifetimes (**k**) of RPpric and RPitc states from zapA-WT and zapA-MT, respectively. The DNA unwinding levels of RPpric and RPitc states were analyzed by averaging over RPpric states (n = 108) and RPitc states (n = 122), and fitted to a single Gaussian function. The lifetimes of RPpric and RPitc states were analyzed and fitted to a single exponential function. **l** Proposed model depicting the impact of productive reiterative initiation on transcription bubble size and gene transcription stimulation. The states of the ternary complexes during regular transcription initiation (left) and productive reiterative initiation (right) are show. The clash between the nascent RNA 5’ end and σR3.2 is indicated by an asterisk.

To test this hypothesis, we compared the DNA sequences of promoters exhibiting “high slippage” and “medium slippage” with those harboring pause sites 10–15-bp downstream of the TSS, as identified by RNET-seq. The notable similarity between these two groups, except for a 1-bp downstream-shifted TIR, elucidated the occurrence of promoter-proximal pausing in promoters featuring productive reiterative initiation (Fig. 5f). Subsequently, we calculated the normalized read count originating from each TSS and observed a gradual increase in read counts with escalating slippage ratios (Fig. 5g). This finding indicates the pivotal role of productive reiterative initiation in enhancing transcription initiation. To elucidate the effect of productive reiterative initiation on transcription initiation, we measured the process of transcription bubble expansion during productive reiterative initiation using single-molecule magnetic trapping (Fig. 5h and i). Interestingly, we found a decrease in the mean transcription bubble size (15.0 ± 0.4 bp, SEM) and an increased lifetime (58 ± 7 s, SEM) of the productive reiterative initiation state compared to the regular transcription initiation state at the mutated *zapA* promoter (18.3 ± 0.5 bp, 39 ± 4 s, SEM) (Fig. 5j and k). We presume that the additional nucleotides added during productive reiterative initiation may not directly influence transcription activity but instead enhance the stability of the initiation complex by reducing the extent of DNA scrunching, thereby counteracting the negative impact of scrunching on complex stability. This stabilization likely provides a longer time-window to facilitate efficient clashes between the nascent RNA 5’ end and σR3.2, ultimately triggering promoter escape (Fig. 5l, Supplementary Fig. 13).

### Role of productive reiterative initiation in regulating cell wall synthesis

To further investigate the function of productive reiterative initiation in gene regulation, we examined its direct impact on gene transcription under different cellular conditions. First, we investigated the correlation between gene transcription changes, as determined by RNA-seq, and variations in slippage ratios. Normalized read counts from RNA-seq within 301-bp DNA regions located 200-bp downstream of TSSs, showing productive reiterative initiation, were compared between the log and stationary phases. The measured Fragments Per Kilobase Million (FPKM) for these DNA regions were significantly higher in the phase with a higher slippage ratio for the corresponding promoters (Fig. 6a). To account for potential transcription associated with non-gene regions, we further examined transcription levels within open reading frames whose start codons were situated within 300-bp from the TSSs. The Transcripts Per Million (TPM) values for the ORFs were consistently higher for promoters exhibiting elevated slippage ratios in each phase (Fig. 6b). Additionally, inhibiting productive reiterative initiation by mutating the transcription initiation regions of the *ffh* and *zapA* promoters significantly reduced the expression of the reporter gene (Fig. 6c), providing further evidence for the role of productive reiterative initiation in enhancing gene transcription.

**Fig. 6.**
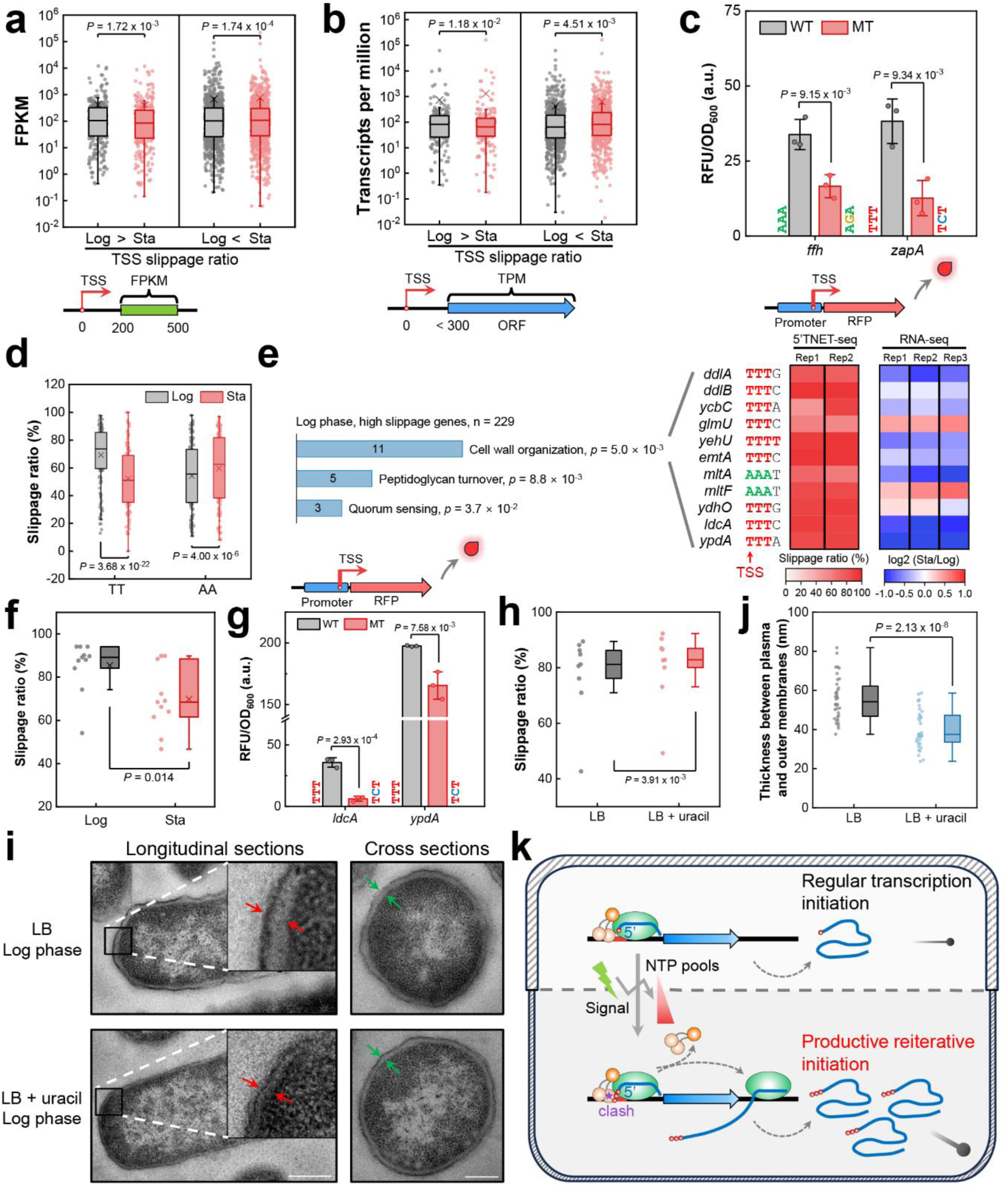
Regulation of cell wall synthesis by productive reiterative initiation. **a** Boxplot illustrating the comparative analysis of transcription levels (FPKM) within DNA regions located 200–500-bp downstream of the TSSs at log and stationary phases. The TSSs exhibiting a slippage ratio 5% higher (n = 275) or lower (n = 832) at the log phase than the stationary phase were applied for analysis. Statistical significance was assessed using the two-tailed Wilcoxon Signed Rank test. **b** Boxplot depicting the evaluation of the transcription level (TPM) of open reading frames positioned less than 300-bp downstream of the TSSs at log and stationary phases. The TSSs with a slippage ratio exceeding 1% (n = 170) or falling below 1% (n = 714) at log phase than stationary phase were used for analysis. *P* values were computed through a two-tailed Wilcoxon Signed Rank test. **c, g** Comparison of promoter strength between wild-type and mutated *ffh*, *zapA* (**c**), *ldcA*, *ypdA* promoters (**g**) using a fluorescent reporter. The trinucleotides adjacent to columns represent the transcription initiation regions of wild-type and mutated promoters. Statistical analysis was performed using a two-tailed student *t*-test (n = 3). **d** Boxplot representation of the slippage ratio alteration from the log phase to the stationary phase for transcription initiation regions commencing with either “TT” (n = 144) or “AA” (n = 136). Statistical analysis employed a two-tailed Wilcoxon Signed Rank test. **e** Gene Ontology and RNA-seq analysis of high slippage genes during the log phase. Heatmaps depict the slippage ratio at the log phase and the log_2_ fold change of 11 genes related to cell wall synthesis. **f** Boxplot illustrating the alteration in the slippage ratio from the log phase to the stationary phase. The *P*-value was statistically calculated via a two-tailed Wilcoxon Signed Rank test (n = 11). **h** Boxplot displaying slippage ratios of promoters associated with cell wall organization in the presence and absence of 1 mM uracil. The two-tailed Wilcoxon Signed Rank test was applied for statistical analysis (n = 9). **i** Visualization of cell wall structures in strains with and without uracil treatment using TEM. Enlarged cell walls images are presented in the insets. Red and green arrows indicate the thickness between the plasma and outer membranes in longitudinal and cross sections, respectively. **j** Boxplot comparing the membrane thickness between the plasma and outer membranes with and without uracil treatment. Statistical analysis was performed using a two-tailed student *t*-test (n = 36). **k** Proposed model illustrating gene transcription regulation through productive reiterative initiation and its impact on cell wall regulation. One red circle represents the 5’ end of RNA, while three red circles represent the addition of two nucleotides at the RNA 5’ end through productive reiterative initiation. The red triangle represents the change in intracellular NTP pools’ concentration.

While non-productive reiterative transcription has been extensively studied at the *pyrBI*, *codBA*, and *upp* promoters for gene regulation, our analysis of 5’TNET-seq data also identified many reads containing homopolymeric tracts, characteristic of transcripts from non-productive reiterative initiation. These reads may associate with non-productive reiterative transcription revealed substantially lower total read counts, 36-fold and 51-fold lower, compared to productive reiterative initiation during the log and stationary phase, respectively (Supplementary Fig. 14a). This discrepancy suggests the prevalence of productive reiterative initiation as a more widespread mechanism compared to non-productive reiterative initiation. Notably, large differences in TSS locations and slippage ratios between the log and stationary phases were observed (Supplementary Fig. 14b and c). Specifically, promoters with “TT” and “AA” TIRs exhibited significant yet opposite changes in slippage ratios from the log phase to the stationary phase, potentially due to changes of NTP concentrations upon phase transition (Fig. 6d).

To investigate the biological significance of productive reiterative initiation, genes with high slippage ratios underwent Gene Ontology (GO) analysis. The analysis indicated that the most significantly enriched group was related to cell wall organization including 11 genes (Fig. 6e, left). Intriguingly, 9 out of the 11 genes possessed a “TTT” TIR, and the majority exhibited reduced transcription during the stationary phase (Fig. 6e, right). Consistent with these findings, the slippage ratios of the 11 corresponding promoters significantly decreased in the stationary phase (Fig. 6f). Furthermore, mutating the TIR to sequences less favorable for productive reiterative initiation significantly reduced the activity of the *ldcA* and *ypdA* promoters (Fig. 6g), suggesting that reduced transcription of cell wall organization genes is mediated by a decline in their productive reiterative initiation. Notably, these 11 genes and their corresponding transcription initiation regions were widespread and highly conserved across 13 bacterial species, especially all of which are pathogens, indicating the conservation of this mechanism in other bacteria (Supplementary Fig. 15).

The correlation between cell wall synthesis and productive reiterative initiation was further explored by culturing cells in LB medium with the addition of 1mM uracil, which induces pyrimidine excess^2^ and promotes productive reiterative initiation. The addition of 1 mM uracil significantly increased the slippage ratios of these genes related to cell wall organization (Fig. 6h). Additionally, analysis of transmission electron microscopy (TEM) images revealed a narrowing of the space between the plasma and outer membranes upon the addition of extra uracil compared to regular log phase cells (Fig. 6i and j). These findings demonstrated dynamic adjustment of productive reiterative initiation in response to environmental signals, such as intracellular NTP concentrations. Elevated productive reiterative initiation, observed in the log phase or with uracil supplementation, enhances the transcription of genes related to cell wall organization, consequently reducing the thickness between the plasma and outer membranes (Fig. 6k).

## DISCUSSION

Reiterative initiation, particularly non-productive reiterative initiation, has been extensively investigated in vitro to elucidate the mechanisms and functions regulating the expression of pyrimidine biosynthetic genes in bacteria^3,4^. Conversely, the existence of productive reiterative initiation in vivo remains inadequately supported by evidence due to the challenging task of detecting and discerning the potential function of extra nucleotides added to the 5’ end of nascent RNA. In addressing this gap, our development of 5’TNET-seq serves as a high-throughput platform to investigate productive reiterative initiation in vivo. Through 5’TNET-seq, we observed a prevalent occurrence of productive reiterative initiation at a genome-wide scale, with over 51% of identified *E. coli* promoters synthesizing slipped RNA. This finding suggests that productive reiterative initiation might represent another universal feature of transcription initiation, along with abortive initiation and promoter-proximal pausing^33,36,37^. Our in vitro transcription data, utilizing low concentrations of NTP, displayed degrees of slippage comparable to those observed under physiological NTP conditions in both in vitro transcription and in vivo 5’TNET-seq data. This observation suggests that high NTP concentration is not essential for productive reiterative initiation, in contrast to non-productive reiterative initiation.

Comparatively, our analysis revealed that dRNA-seq and SEnd-seq data, capable of detecting TSSs, also show mismatched nucleotides at the 5’ end of transcripts originating from certain promoters^31,32^, indicative of productive reiterative initiation. However, 5’TNET-seq excels in detecting a greater number of productive reiterative initiation events and obscure TSSs. Leveraging the enrichment of nascent RNA with slippage through the pull-down of native elongating ternary complexes, 5’TNET-seq outperforms other methods in detecting productive reiterative initiation, which employ total RNA and have low sensitivity for detecting unprocessed RNA. This enrichment, facilitated by the high frequency of transcription pausing^38,39^, enhances the resolution for identifying less conserved promoters. Moreover, 5’TNET-seq provides a more efficient and expeditious means to quantify the slippage ratio at a genome-wide level compared to traditional reiterative transcription determination methods involving sequencing gel electrophoresis analysis^2,26^.

Metagene analysis of 5’TNET-seq data reveals that homopolymeric tract in the transcription initiation region, especially A-tract or T-tract, is a key element for productive reiterative initiation. In addition, these homopolymeric tracts contribute to all transcript slippages during the entire transcription process, including initiation, elongation, and termination^22,24,34,40^. Notably, the *pyrG* promoter in *E. coli*, with a “CCC” TIR, did not exhibit a high slippage ratio (10.1%) of productive reiterative initiation, as extensively studied in *B. subtilis* with a G-tract transcription initiation region. This concurs with the notion that weaker hydrogen bonds between DNA and nascent RNA base pairs, as seen in the stabilization hierarchy (dCrG > dGrC > dTrA > dArU)^41^, facilitate RNA slippage. Homopolymeric DNA sequences containing at least nine “A” or “T” nucleotides are favored for transcript slippage during elongation^42,43^, while “AA” and “TT” dinucleotides are capable of stimulating slippage during transcription initiation. This suggests that a less stable short RNA-DNA hybrid induces genome-wide productive reiterative initiation.

Moreover, our findings indicate that productive reiterative initiation is not solely characterized by the addition of a single nucleotide per cycle; up to four nucleotides can be added in a single slippage cycle at over 100 promoters as identified by 5’TNET-seq. Longer spacer lengths and the presence of matched nucleotides with the transcription initiation region in these promoters imply that a larger space between the –10 region and the TSS is necessary to accommodate slipped nucleotides, and the matched nucleotides allow base pairing of the RNA 5’ end to DNA after slippage, contributing to the stabilization of the slipped nascent RNA. These findings are reminiscent of the *rrnB* P1 and λP_L_ promoters, where the nascent transcript slips back three nucleotides to an alternative stable state^12,44^. The weakest dA:rU base pairing at T-tract transcription initiation regions, providing matched nucleotides upstream for RNA-DNA base pairing after slippage, contributes to high slippage ratios and more slippage cycles at promoters with homopolymric T-tract.

Non-productive reiterative initiation regulates gene transcription by aborting transcripts to attenuate transcription initiation^3–5^. In contrast, productive reiterative initiation switches back to regular transcription after diverse rounds of transcript slippage, and its impact on transcriptional regulation requires elucidation. Unlike transcript slippage during transcription elongation, which typically occurs in ORFs, such as slippage in insertion sequence genes^42,45^ that interrupts or rescues gene function by introducing frameshifts^10,43^, transcripts from productive reiterative initiation that adds extra nucleotides to the RNA 5’ end generally do not generate frameshifts or influence translation. In regular transcription initiation, RNA synthesis starts at TSS by RNAP, generating a perfectly base-paired RNA-DNA hybrid until the hybrid exceeds 9–10-bp^46,47^. At this point, the RNA 5’ end enters the RNA-exit channel in RNAP preventing RNA base pairing with the template DNA^48^. In this case, the mismatched nucleotides at the 5’ end of transcripts from productive reiterative initiation behave similarly with those from regular transcription. This suggests that the mismatched 5’ end in the longer nascent RNA functions before the formation of a 9–10-bp RNA-DNA hybrid.

Transcription initiation involves the reversible scrunching of promoter DNA leading to increase the transcription bubble size, followed by either promoter escape or a return to the open promoter state. The σR3.2 region, partially located in the RNA exit channel, is displaced by the extending nascent RNA’s 5’ end before promoter escape^49,50^. Although the extra nucleotides added by productive reiterative initiation do not alter the frames of the ORFs, productive reiterative initiation ultimately shortens the transcription bubble size and the RNA-DNA hybrid without reducing the RNA length. This shortening may alleviate steric hindrance within the transcription bubble and reduce extensive scrunching, resulting in a stabilized reiterative transcription initiation complex and a steady progressive clash between the 5’ end of nascent RNA and σR3.2 through direct collision. This process is likely to enhance promoter escape, positively regulating transcription initiation by increasing transcription efficiency compared to regular transcription initiation, which may be more prone to relaxing the initiation complex to an open promoter state and aborting transcripts. This conclusion aligns with previous mutation analyses on a *lac* +10A promoter mutant and *pyrG* promoter, demonstrating that a single nucleotide deletion or mutation in the homopolymeric tract of the transcription initiation region significantly reduces promoter activity^14,51,52^. Similarly, our single mutation from “T” to “A” on the *zapA* promoter also almost abolished run-off transcription (Fig. 5d), emphasizing the critical role of productive reiterative initiation in gene transcription. Therefore, productive reiterative initiation constitutes another transcription process enhancing initiation through a novel mechanism, independent of transcription activators. In circumventing the need for protein regulators^53,54^, productive reiterative initiation offers a direct and prompt mechanism to regulate cell wall synthesis in response to environmental signals, such as NTP concentrations influenced by nutrient sources and cell growth phases^55^.

While 5’TNET-seq provides a comprehensive and quantitative approach to investigate productive reiterative initiation, it is important to note that non-productive reiterative initiation, generating heterogenous lengths of polynucleotide tracts, could also be studied using this method. The sequencing reads reveal that the count of reads from productive reiterative initiation is an order of magnitude higher than that from non-productive reiterative initiation, indicating the more extensive and widespread occurrence of productive reiterative initiation. However, due to the principle of the pipeline, which extracts reads with slippage based on mismatched nucleotides, reads generated by productive reiterative initiation and precisely aligned to the genome are not extracted and analyzed. This suggests that the actual events and modes of productive reiterative initiation may surpass the identified cases presented here. Further studies are warranted to investigate whether regulators influencing the stability of the RNA-DNA hybrid or occupying the RNA exit channel (e.g., Rho, NusA) could regulate productive reiterative initiation.

## METHODS

### *E. coli* strain and growth conditions

*E. coli* strain β’-WT (W3110 *rpoC*-6×His::*kan*) constructed in the previous study^56^ was cultivated in Luria-Bertani (LB) medium (10 g L^−1^ tryptone, 5 g L^−1^ yeast extract, 10 g L^−1^ NaCl) or on LB plates supplemented with 50 μg L^−1^ kanamycin at 37°C. An overnight culture of cells was appropriately diluted to 250 ml of fresh LB medium, initiating with an OD_600_ of 0.02, and subsequently incubated at 37°C. Cells grown to mid-log phase (OD_600_ = 0.5) and stationary phase (OD_600_ = 2.0) were harvested for further analyses.

### 5’TNET-seq library generation

Cell collection, lysis, and native elongating complex purification. *E. coli* cells, cultivated to log and stationary phases, were rapidly chilled by mixing with an equal volume of frozen crush buffer (20 mM Tris-HCl pH 7.8, 10 mM ethylenediaminetetraacetic acid (EDTA), 100 mM NaCl, 1 M Urea, 25 mM NaN_3_, 2 mM β-mercaptoethanol, 10% ethanol, 0.4% NP40, 1 mM PMSF). The cells were subjected to centrifugation, flash-frozen in liquid nitrogen, and stored at –80°C until cell lysis. Upon thawing, the cells were resuspended in lysis buffer (10 mM Tris-HCl pH 7.8, 1 mM EDTA, 1mM spermine, 0.1% Triton X-100, 100 mM NaCl, 4 mM MgCl_2_, 4 mM CaCl_2_, 2.5% PEG8000, 100 μg ml^−1^ bovine serum albumin (BSA), 1 × protease inhibitors) and lysed using 120 kU Ready-Lyse lysozyme (Lucigen). Native elongating complexes were released from chromosome by treatment with 6 U Turbo DNase (Invitrogen) and 100 U DNase I (Roche) at 23°C for 10 min. The released elongating complexes were pulled down using 200 μl Ni-NTA agarose beads by incubation at 4°C for 10 min with shaking. The beads, housing the native elongating complexes, underwent three washes with 1 ml of cold wash buffer (20 mM Tris-HCl pH 7.8, 1 M betaine, 5% glycerol, 2 mM β-mercaptoethanol, 2.5 mM imidazole). The beads were then loaded onto 0.5 ml Ultrafree-MC centrifugal filters (Millipore). Finally, the elongating complex was eluted using wash buffer containing 0.3 M imidazole.

Nascent RNA extraction, size fractionation, and adapter ligation. Nucleic acids from the elongating complexes were extracted and purified using phenol:chloroform:isoamylalcohol (PCI; 25:24:1) and chloroform. Isopropanol precipitation was performed to obtain the nucleic acid pellet, which was then washed by cold 80% ethanol, air dried and dissolved in 12 µl nuclease-free water. Genomic DNA contamination was eliminated by treating with 13 µl DNase mixture (2 U Turbo DNase, 10 U DNase I and 10 U SuperaseIn RNase inhibitor) at 37°C for 15 min. The remaining nascent RNA was further purified, precipitated, and fractionated on a 15% TBE-Urea gel. The gel containing the nascent RNA (8–100-nt) was excised, crushed, and extracted by 200 µl nuclease-free water at 70°C for 15 min with shaking. The gel-extracted nascent RNA in 5 µl nuclease-free water was ligated to 10.7 pmol barcode DNA linker (Supplementary Table 2) using 200 U truncated T4 RNA ligase 2 (NEB) overnight at 16°C. RNA-DNA chimeras generated was then extracted, precipitated, and subjected to treatment with 5’-phosphate-dependent exonuclease (Epicentre) as required.

TEX treatment, reverse transcription, and circularization. The RNA-DNA chimeras were denatured at 70°C for 2 min, and subsequently treated with or without 2 U 5’-phosphate-dependent exonuclease (Epicentre) at 30°C for 1h. Following PCI, chloroform extraction and isopropanol precipitation, the RNA-DNA chimera was reverse transcribed by the addition of 3 µM phosphorylated reverse transcription primer (Supplementary Table 2) in 1 × PrimeScript buffer containing 0.5 mM dNTPs, 5 mM DTT, 0.6 U µl^-1^ SuperaseIn RNase inhibitor, and 10 U µl^-1^ PrimeScript Reverse Transcriptase (Takara), followed by incubation at 48°C for 30 min. Subsequently, 2 U RNase H (NEB) was added to hydrolyze the RNA at 37°C for 15 min, and the resulting cDNA was fractionated on a 10% TBE-Urea gel. The cDNA products within the 80–175-nt range were then extracted, precipitated, and dissolved in 4 µl of nuclease-free water. Subsequent cDNA circularization was performed at 60°C for 4 h, utilizing 40 U of ssDNA Ligase and 50 µM ATP (Lucigen).

DNA library generation and Illumina sequencing. The circularized DNA was employed as templates for PCR amplification of the sequencing libraries, using Illumina index primers and PrimeSTAR Max DNA polymerase. The resulting PCR product was subjected to electrophoresis on an 8% TBE gel, and the target DNA product, ranging approximately from 150 to 240-bp, was extracted using a DNA soaking buffer (0.3 M NaCl, 10 mM of Tris-HCl, pH 8.0, 0.97 mM of EDTA) after overnight shaking. Following precipitated with isopropanol, the DNA libraries were washed with 180 µl of cold 80% ethanol and ultimately dissolved in 8 µl TE buffer (10 mM Tris-HCl, pH 8.0). The size and concentration of the DNA libraries were assessed using an Agilent 2100 bioanalyzer and quantified via qRT-PCR before sequencing. The final DNA libraries were pooled and sequenced on an Illumina NovaSeq 6000 platform (2 × 50 bp paired end) in SP mode.

### 5’TNET-seq data processing pipeline

Read mapping, TSS and PS identification. The raw sequencing data was demultiplexed and Illumina adapters at the 3’ end of R1 and R2 reads were trimmed by Cutadapt^57^. Subsequent merging of R2 and R1 reads with FLASH^58^ was followed by the removal of PCR duplicates using Clumpify from the BBMap suite. Additionally, a 6-nt random barcode at the 3’ end was trimmed by Cutadapt. The resultant reads were aligned to the *E. coli* genome NC_000913.2 through Bowtie^59^, allowing a maximum of two mismatches in a single read. Uniquely aligned reads and unmapped reads were extracted for further analysis, while multi-mapped reads were discarded. The 5’ end and 3’ end coordinates of all uniquely mapped alignments at each genomic position (C5 and C3) were recorded by BEDTools^60^ to identify the original TSSs and pause sites. A genomic position was identified as a transcription start site if the C5 count equal to or greater than 5 reads per million, with C5 being at least 20-fold higher than the median C5 within a 51-nt window size and exhibiting a 2-fold increase compared to the adjacent upstream position. A genomic position was categorized as a transcription pause site if the C3 count was equal to or greater than 10 reads per million, with C3 being at least 20-fold higher than the median C3 within a 51-nt window size.

Slippage ratio, pattern, and nucleotide addition analysis. Unmapped reads were aligned to the genome using Bowtie after trimming 5’ end nucleotides. In each cycle, read sequence was divided into two parts: the 5’ end nucleotide (slippage part) and the read sequence after trimming (residual part). The latter, with one nucleotide trimmed from the 5’ end by Cutadapt, was mapped to the genome using the Bowtie aligner. Multiple mapping reads were discarded, and the unique alignment with the corresponding trimmed 5’ end nucleotide was used for slippage analysis. Unmapped residual parts were processed in the next cycle, and the trimmed slippage parts were combined until the reads were aligned to unique or multiple genomic locations. Reads with mismatches in the first three positions (mismatch part) from the 5’ end, which uniquely aligned before or after trimming, were extracted for slippage analysis using custom scripts. The first nucleotide downstream of the last mismatched nucleotide, matching the 5’ end nucleotide, was redefined as the 5’ end of the read. The cumulative length of the slippage part and mismatch part was considered the number of nucleotides added by productive reiterative initiation. The 5’ end and 3’ end read coordinates for all unique alignments were updated to correct TSSs and PSs. The slippage ratio (%) for each TSS was calculated by dividing the slippage read counts by the total read counts derived from the corresponding TSS. Slippage patterns were classified by comparing the mismatch part and slippage part of the reads with the transcription initiation region. If the first four, three or two nucleotides of the slippage read were not the same as the initial nucleotide and exactly matched the TIR, the slippage pattern was sequentially classified as four (4nc), three (3nc) or two (2nc) nucleotides slippage per cycle of productive reiterative initiation. If only one slipped nucleotide or the first two nucleotides added matched the TSS, the slippage pattern was classified as one (1nc) nucleotide slippage. Otherwise, a promoter was designated as 0nc, representing aberrant nucleotide addition.

### 5’ RACE and data processing

The procedure for extracting nascent RNA from cells collected at log phase (OD_600_ = 0.5) for 5’ RACE was identical with that of 5’TNET-seq. Following genomic DNA digestion and TEX treatment, nascent RNA was extracted using PCI and chloroform, and then recovered through isopropanol precipitation. The RNA product, dissolved in 10 µl nuclease-free water, was treated with 10 U RNA 5’ pyrophosphohydrolase (NEB) at 37°C for 30 min. The reaction was halted by adding 1 µl 0.5 M EDTA. The resulting RNA product was purified by PCI and chloroform extraction, precipitated, and solubilized in 5 µl nuclease-free water. Following denaturation at 70°C for 2 min, a 15 µl ligation mixture containing 100 pmol of the 5’ adapter and 10 U T4 RNA ligase 1 was mixed the RNA product and incubated overnight at 37°C with shaking. The ligation product was purified and reversed transcribed using 25 pmol pooled primers (IDT, Supplementary Table 2) for target TSSs at 48°C for 30 min. The reverse transcription product was treated through incubation with 2 U RNase H and separated by electrophoresis on a 10% TBE-Urea gel. The cDNA within the 65–80-nt range was recovered and used as the DNA templates for PCR amplification of the sequencing libraries. The concentration of 5’ RACE libraries were quantified by the Agilent 2100 bioanalyzer and sequenced on Illumina MiSeq with 2 × 150 bp paired-end reads. The reads were processed identically to 5’TNET-seq, and a position was defined as a transcription start site when its 5’ end read counts was at least 20-fold higher than the median read count within a surrounding 51-nt window size and not less than 10 reads per million.

### In vitro transcription assays

Gel electrophoreses. DNA templates, encompassing the *ffh* and *zapA* promoters within the range of –100 to +40-bp relative to the TSSs, were PCR amplified from the *E. coli* MG1655 genome. Mutant DNA was generated by annealing and extending primers with specific point mutations. In vitro transcription reactions were performed by adding 20 nM DNA template and 50 nM RNAP holoenzyme in transcription buffer (40 mM Tris-HCl pH 8.0, 1 mM dithiothreitol, 0.1 mg ml^-1^ BSA, 10 mM MgCl_2_, 50 mM KCl). When indicated, 50 nM GreB was added to the reaction mixture. After a 10-min incubation at 37°C to form the open promoter complex, reactions were initiated by adding different sets of NTPs: 20 µM UTP, CTP, GTP, and either 20 or 100 µM ATP for the *ffh* promoter; 20 µM ATP, CTP, GTP, and either 20 or 100 µM UTP for the *zapA* promoter. RNA transcripts from the *ffh* and *zapA* promoters were labeled with 2– 10 µCi of [α-^32^P] ATP or [α-^32^P] UTP (PerkinElmer), respectively. Reactions were carried out at 37°C for 10 min and terminated by adding an equal volume of stop buffer (10 M Urea, 250 mM EDTA pH 8.0, 0.05% xylene cyanol and bromphenol blue). The reaction products were electrophoresed on a 23% (10:1, acrylamide:bisacrylamide) polyacrylamide gel with 7 M urea and visualized using the GE Typhoon TRIO Imager (GE Healthcare).

High-throughput sequencing. DNA sequences from the –60 to +39-bp region relative to the TSSs of target promoters were applied for sequencing in vitro transcription products. Single-stranded DNA sequences appended with a 20-nt unique sequence at the 3’ end (Supplementary Table 2) were synthesized in an oligo pool (IDT). Pooled oligos were extended and converted to double-stranded DNA using a universal primer annealed to the 3’ end unique sequence, followed by gel purification after separation on an 8% TBE gel. In a 20 µl reaction mixture, 100 nM pooled DNA template and 250 nM RNAP holoenzyme were used to generate the open promoter complex after incubation at 37°C for 10 min. Then, three different concentrations of NTPs—low UTP (200 µM ATP, CTP, GTP and 40 µM UTP), low ATP (200 µM UTP, CTP, GTP and 40 µM ATP) and regular NTPs (200 µM ATP, UTP, CTP and GTP)—were added to initiate transcription at 37°C for 10 min. Sequencing libraries were constructed using transcription products similar to 5’TNET-seq and subjected to paired-end sequenced (2 × 150 bp) on an Illumina MiSeq platform. The sequencing data were analyzed as employed for 5’TNET-seq.

### Single-molecule magnetic trapping assay and data analysis

DNA fragments containing the wildtype (C_-1_T_+1_T_+2_T_+3_) and mutant (C_-1_T_+1_A_+2_T_+3_) *zapA* promoter region (–60 to +20-bp relative to TSS), respectively, followed by a 77-bp transcription unit and the *E. coli his* terminator (Supplementary Table 2) were synthesized and subcloned into pUC18 through the flanked KpnI sites. DNA constructs for single-molecule magnetic trapping assay were prepared by digesting the obtained plasmids with XbaI and SbfI (New England Biolabs), and then ligated with two 1-kb DNA fragments modified with multiple biotin groups and digoxigenin groups through the XbaI and SbfI sites^61^, respectively. As previously described^62^, glass surfaces were cleaned and sandwiched into a flow chamber and then functionalized with anti-digoxigenin (Roche). DNA constructs were tethered between the glass surface and a magnetic bead (Dynabeads MyOne Streptavidin C1; Life Technologies) and then extended and supercoiled (F = 0.1 pN, 1 pN = 10^-12^ N; superhelical density = 0.023 or +5 for positive supercoiling) via a homemade magnetic trap. Bead position reflecting DNA extension changes was monitored via a BeadTracker software suite developed by N. H. Dekker lab^63^. Single-molecule data were collected at 50 Hz and filtered at 1 Hz. Experiments were performed in transcription buffer (20 mM HEPES K+ pH 8.0, 100 mM KCl, 8 mM MgCl2, 0.5 mg/mL BSA, 0.1% (v/v) Tween 20, 10 μM ZnCl2, 1 mM DTT) at room temperature (around 28°C). Successful escaped transcription events were observed at 2 nM RNAP, 200 µM NTPs and 50 nM GreB, and the lifetime and mean amplitude changes of the initiation state were analyzed via custom Matlab codes as previously^62^.

### Transmission electron microscopy

*E. coli* MG1655 cells cultured in LB medium, with or without the addition of 1 mM uracil, were harvested at the log phase (OD_600_ = 0.5). After collection, cells were washed four times with phosphate-buffered saline (PBS) and fixed in PBS containing 2.5% glutaraldehyde overnight at 4°C. Following fixation, cells were subjected to three washes with PBS, two washes with distilled water, and immersion in 1% osmium tetroxide at 4°C for 1 h. The samples were subsequently washed three time with PBS. Dehydration of the samples were achieved through a series of ethanol solutions with increasing concentrations, each step lasting 10 min. Finally, the processed cells were embedded in LR white resin. Embedded cells were sectioned into ultrathin slices, approximately 70 nm in thickness. The imaging process employed a Hitachi HT7700 high-resolution transmission electron microscope.

### RNA-seq and data analysis

Cells grown to mid-log and stationary phases were promptly quenched by the addition of an equal volume of cold quench buffer (5% acid Phenol:Chloroform in ethanol). The quenched cells were then collected by centrifugation and resuspended in 1 ml TRIzol reagent (Invitrogen). Total RNA extraction was carried out using PCI and chloroform, followed by isopropanol precipitated. The extracted RNA underwent treatment with TURBO DNase and DNase I to eliminate genomic DNA contamination. The resulting RNA was further purified using the RNeasy Mini Kit (Qiagen). RNA was quantified using the Agilent 2100 bioanalyzer and applied to construct sequencing libraries using the TruSeq Stranded Total RNA Library Prep Kit (Illumina). The generated DNA libraries were sequenced at the Center for Cancer Research Sequencing Facility, using the NextSeq 2000 platform with 2 × 100 bp paired-end reads. Raw sequencing reads were mapped to the *E. coli* genome (NC_000913.2) using Bowtie 2^64^. Counting of aligned reads for each gene was conducted using HTSeq^65^, and differential gene expression analyses was executed through DESeq2^66^.

## Data availability

All 5’TNET-seq and RNA-seq data from this study have been deposited to NCBI’s Gene Expression Omnibus (GEO) database under the accession number GSE248682. The dRNA-seq data were obtained from GEO with the accession number GSE55199^31^. Source data are provided with this paper and deposited at Mendeley Data (DOI: 10.17632/t9kp3ntg8n.1).

## Code Availability

The custom scripts utilized in processing 5’TNET-seq data to identify pause sites and calculate slippage ratios are available on GitHub (https://github.com/ZSunTIB/5-TNET-seq).

## Acknowledgements

We thank Prof. Kuanqing Liu for critical reading of this manuscript. We thank the NIH Intramural Sequencing Center and the CCR Sequencing Facility for Illumina sequencing. This work was financially supported by the National Key Research and Development Program of China (2022YFC2106200 to Z.S.), the National Natural Science Foundation of China (32270031 to Z.S.), the Tianjin Synthetic Biotechnology Innovation Capacity Improvement Project (TSBICIP-CXRC-064 to Z.S.), the Hundred Talents Program of the Chinese Academy of Sciences (to Z.S.) and the Intramural Research Program of the National Institutes of Health, National Cancer Institute, Center for Cancer Research (to M.K.).

## Author contributions

Z.S. and M.K. conceived and designed the project. Z.S. developed and performed 5’TNET-seq. Z.S. developed the 5’TNET-seq data processing pipeline. Z.S. conducted 5’RACE, RNA-seq and biochemical experiments. Y.Z and S.W. performed magnetic trapping assay and data analysis. G.Z. performed transmission electron microscopy experiment. Z.S. analyzed sequencing and TEM data. Z.S. wrote the manuscript.

## Additional information

Extended data is available for this paper.

