## Supplementary information for "Nascent RNA profiling reveals regulation of gene transcription through productive reiterative initiation in bacteria"

**TITLE**

[*]Correspondence:

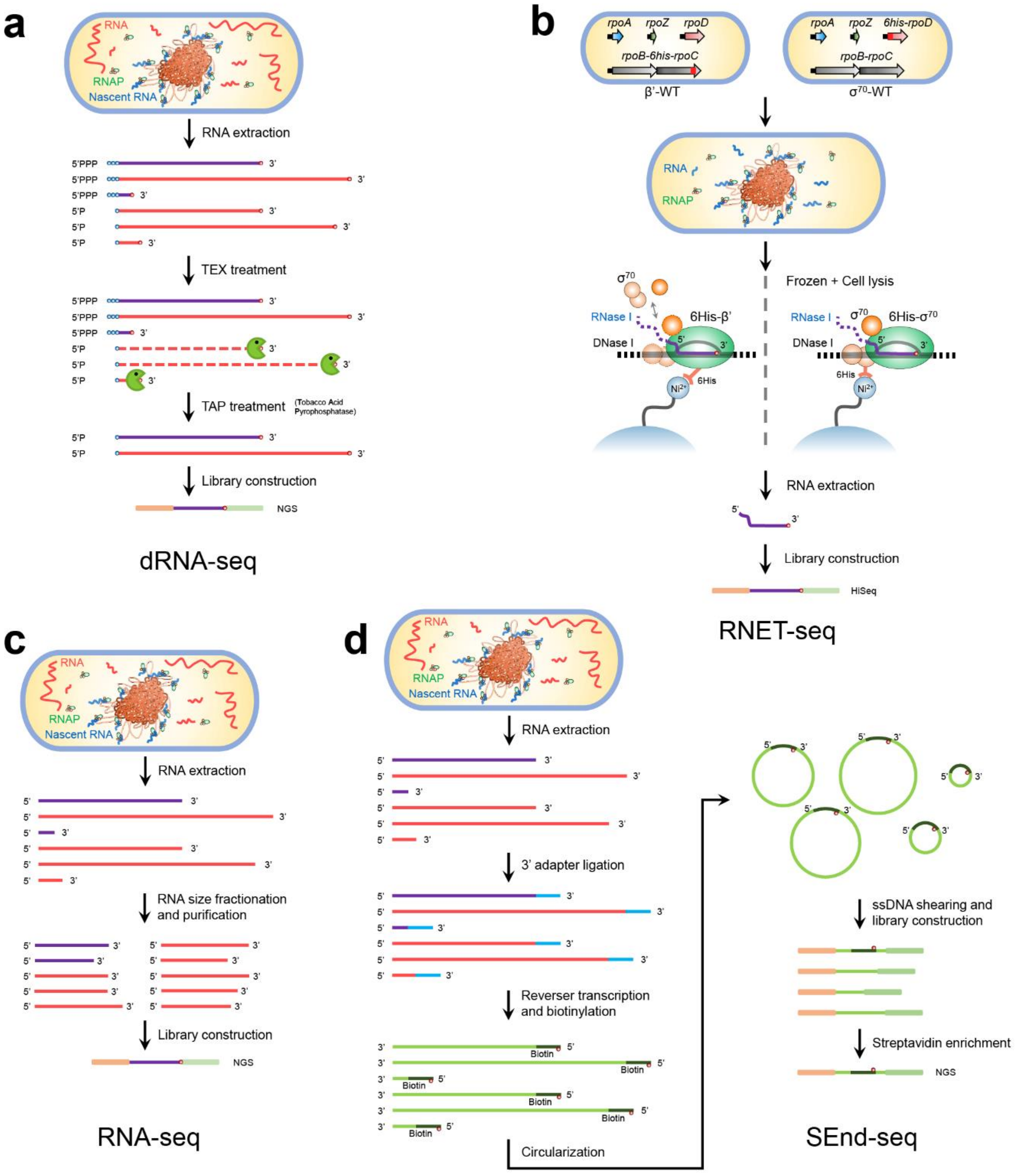


**Supplementary Fig. 1 | Schematic representation of the dRNA-seq, RNET-seq, RNA-seq and SEnd-seq workflows.** The workflows of RNET-seq and SEnd-seq were adapted and modified from previous studies[1,2].

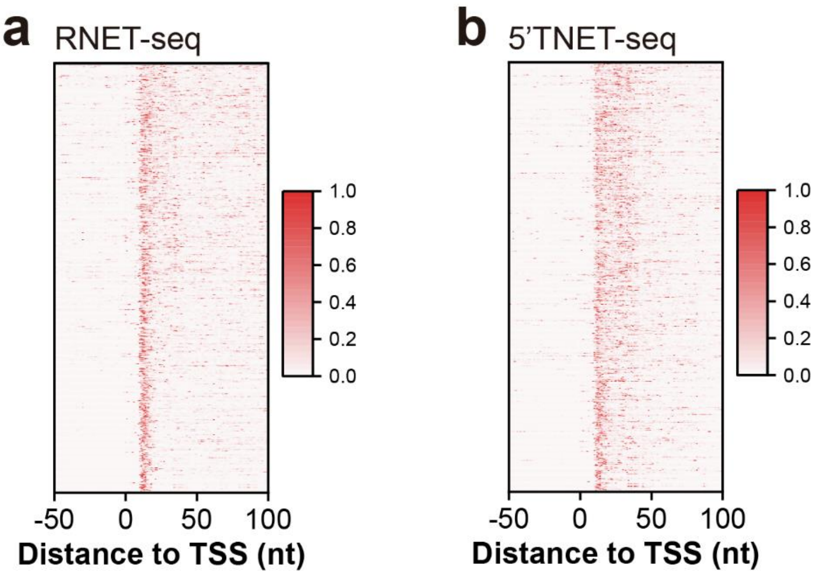


**Supplementary Fig. 2 | Heatmaps illustrating the normalized pause ratios of RNAP in RNET-seq and 5’TNET-seq.** Analyses were conducted using pause sites identified in the promoter-proximal regions (+1 to +40 relative to TSSs) by RNET-seq (n = 607). The maximum read count of the RNA 3’ end within the –50 to +100 region relative to the TSSs was normalized to 1. The resulting normalized pause ratios for each region detected by RNET-seq (**a**) and 5’TNET-seq (**b**) are depicted.

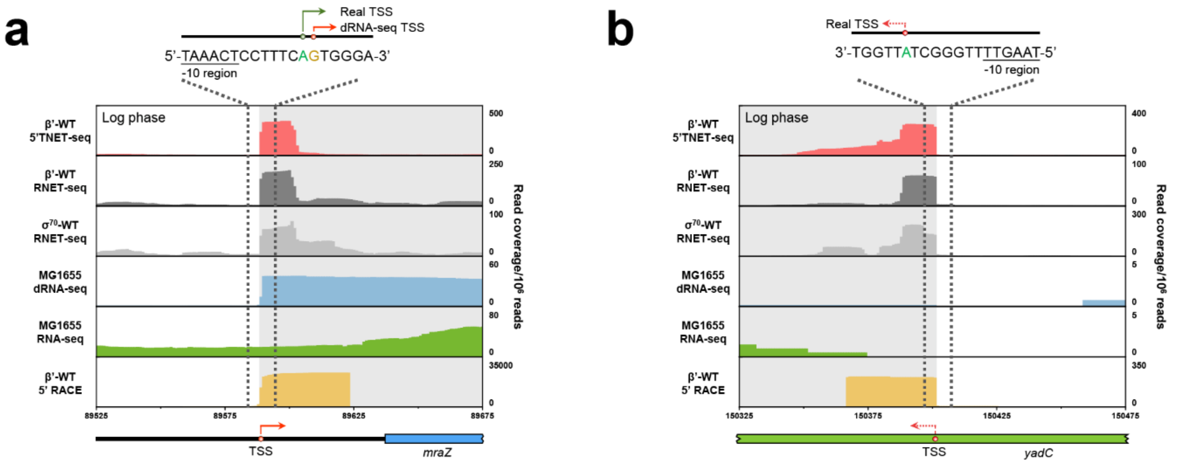


**Supplementary Fig. 3 | Sequencing profiles of DNA regions revealing newly identified TSSs via 5'TNET-seq.** Sequencing profiles of DNA regions displaying novel TSSs located upstream of the *mraZ* gene's ORF (**a**) or within the ORF of the *yadC* gene (**b**). The sequences of the TSS regions and the precise TSS locations are depicted atop the profiles.

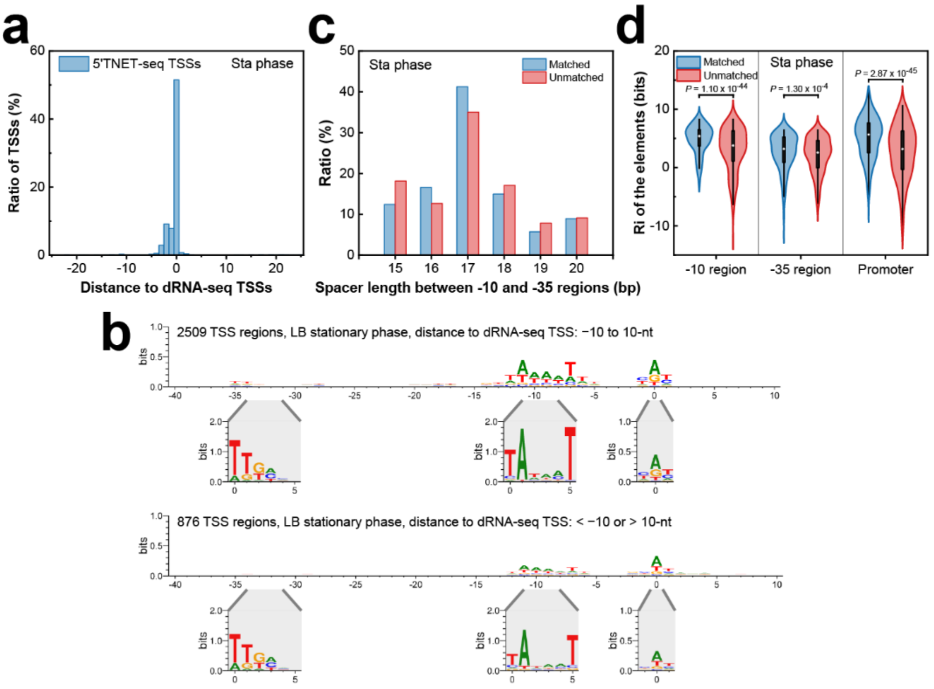


**Supplementary Fig. 4 | Characteristics of TSSs identified by 5'TNET-seq during the stationary phase. a** Histogram displaying the distance between the TSSs identified by 5'TNET-seq during the stationary phase and their nearest dRNA-seq TSSs counterparts. **b** Sequence logos representing the promoter regions identified by 5'TNET-seq, categorized by the proximity of their TSSs to the nearest dRNA-seq TSSs (≤ 10-nt or > 10-nt). Sequence logos for the –35, –10 and TSS regions are presented below following alignment. **c** Comparative analysis of the spacer length for promoters with "matched" or "unmatched" TSSs between 5'TNET-seq and dRNA-seq during the stationary phase. **d** Violin plot showing the Ri of the –10 region, –35 region, and the entire promoter for "matched" (n = 2509) or "unmatched" (n = 876) TSSs between 5'TNET-seq and dRNA-seq during the stationary phase. In this and all subsequent violin plots, the box represents the interquartile range, the whiskers extend to 1.5 times the interquartile range, and the white cycle indicates the median. Statistical significance was assessed using a two-tailed Mann-Whitney *U*-test.

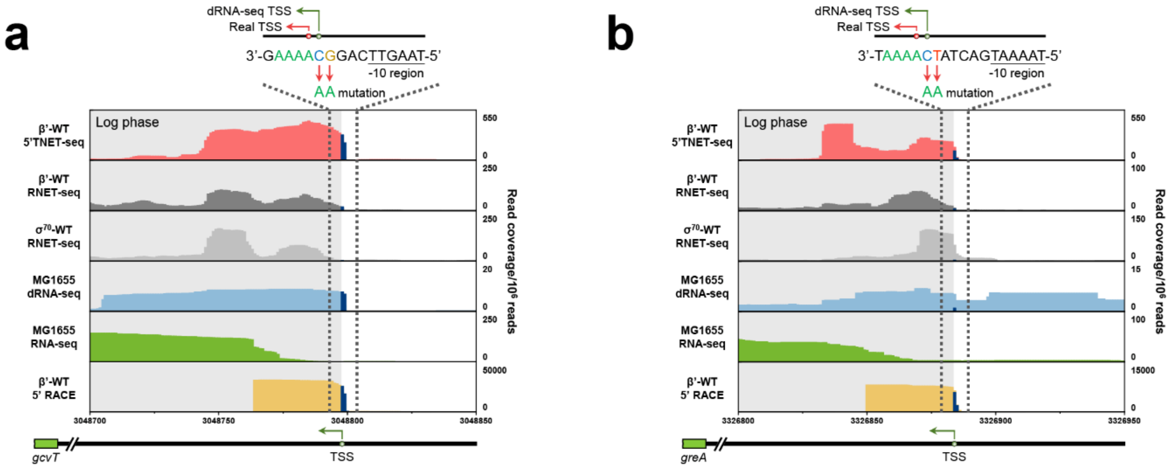


**Supplementary Fig. 5 | Sequencing profiles of DNA regions housing TSSs incorrectly annotated by dRNA-seq due to productive reiterative initiation.** Sequencing profiles displaying TSSs located upstream of the ORFs of the *gcvT* (**a**) and *greA* (**b**) genes. The green arrows denote TSSs that were inaccurately labeled using dRNA-seq. The red arrows indicate the authentic TSSs.

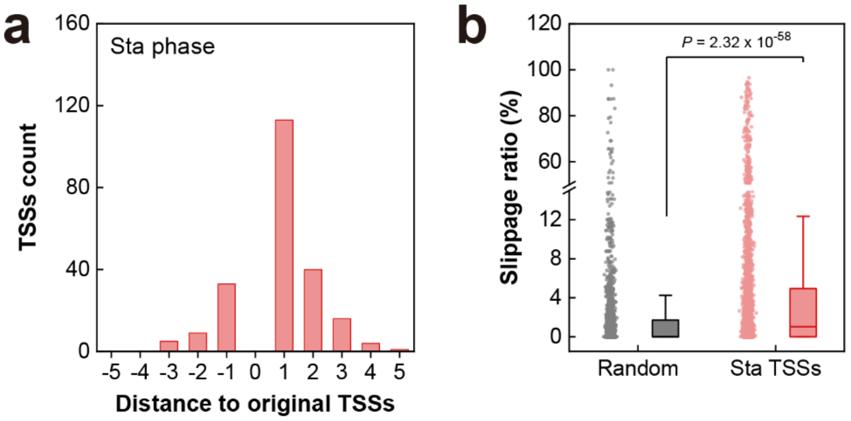


**Supplementary Fig. 6 | Characteristics of TSSs identified by 5'TNET-seq during the stationary phase. a** Histogram illustrating the distribution of distance between TSSs before and after correction, attributed to productive reiterative initiation. **b** Boxplot depicting the slippage ratio of TSSs during the stationary phase, compared to an equivalent number of randomly selected sites (n = 3509). In this and all subsequent boxplots, the box corresponds to the interquartile range, the line denotes the median, and the whiskers extend to 1.5 times the interquartile range. Statistical analysis was conducted using a two-tailed Mann-Whitney *U*-test.

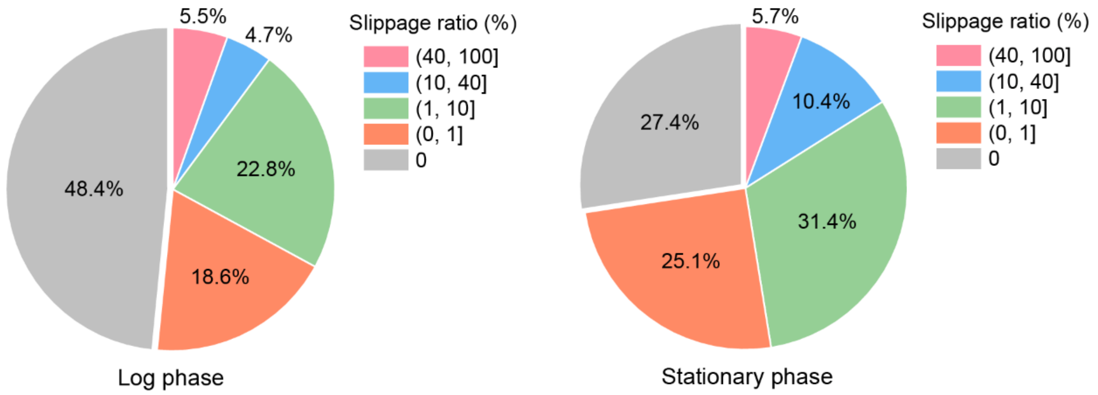


**Supplementary Fig. 7 | Relative proportion of TSSs detected by 5'TNET-seq with different slippage ratios.** Pie charts illustrate the distribution of TSSs identified by 5'TNET-seq at the log (left, n = 4188) and stationary (right, n = 3622) phases.

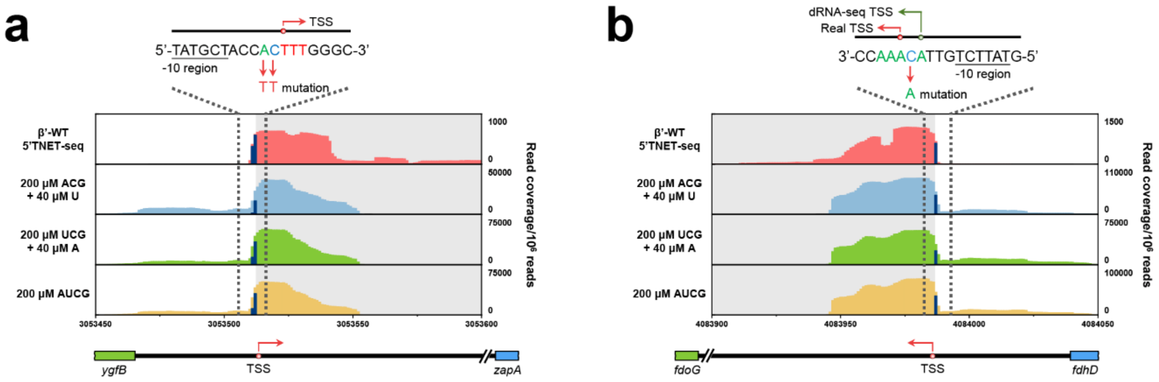


**Supplementary Fig. 8 | In vitro transcription assays confirm the slippage ratios of promoters with “AAA” or “TTT” TIRs.** Profiles of 5’TNET-seq at the log phase, along with in vitro transcription assays, to validate the slippage ratios of the promoters upstream of the ORFs of *zapA* (**a**) and *fdoG* (**b**) genes. The TSS regions are delineated, with the actual TSSs and erroneously labeled TSSs indicated atop the profiles.

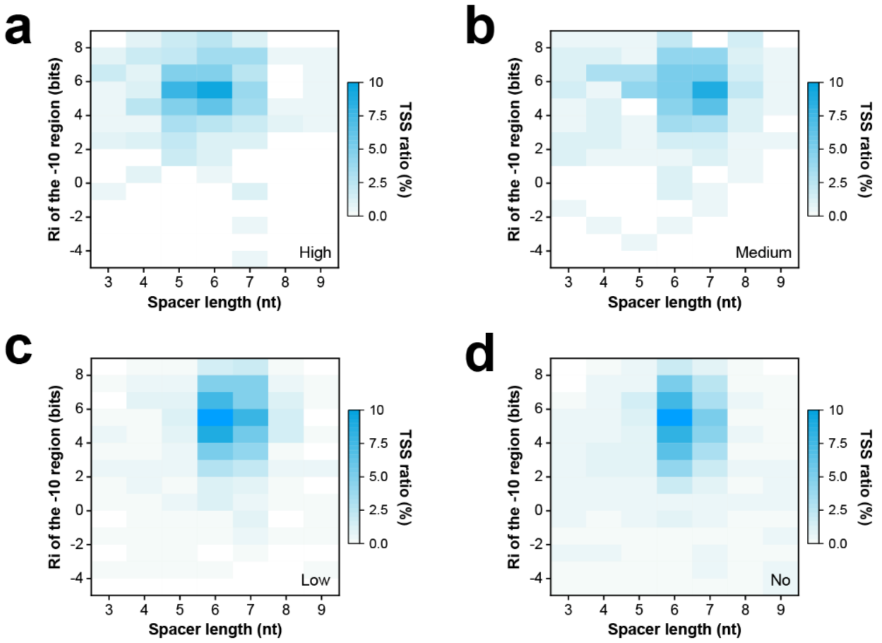


**Supplementary Fig. 9 | Heatmaps showing the distribution of the –10 region Ri and spacer length for TSSs with different slippage ratios.** Heatmaps for the TSSs categorized into "high" (**a**), "medium" (**b**), "low" (**c**), and "no slippage" (**d**) are displayed, respectively.

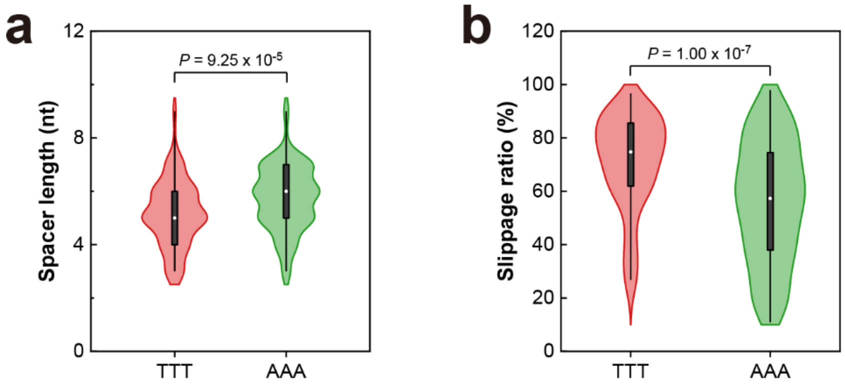


**Supplementary Fig. 10 | Characteristics of promoters with homopolymeric trinucleotide initiation regions.** Violin plots presenting the spacer length (**a**) and slippage ratio (**b**) for promoters initiating with homopolymeric trinucleotides “TTT” (n = 126) or “AAA” (n = 129) relative to the TSSs. Statistical analysis was exclusively performed on promoters exhibiting high and medium slippage using a two-tailed Mann-Whitney *U*-test.

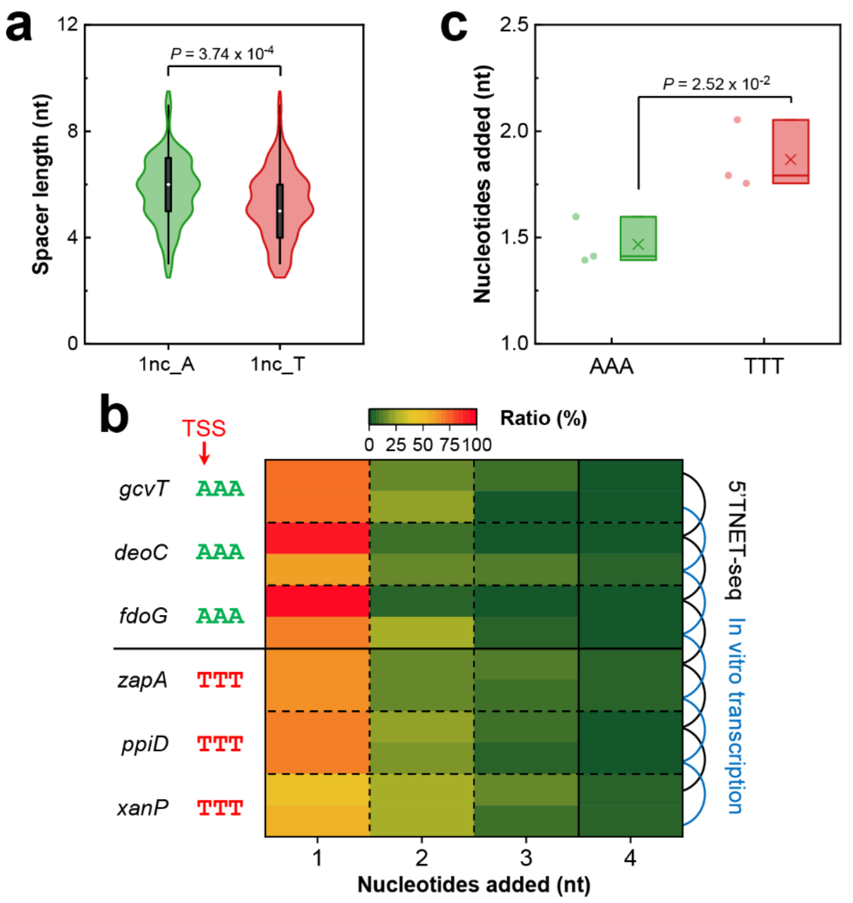


**Supplementary Fig. 11 | Experimental validation of nucleotides addition via productive reiterative initiation. a** Violin plot analysis of spacer length in promoters belonging to 1nc with "A" (n = 136) or "T" (n = 144) additions. Two-tailed Mann-Whitney *U*-test was applied for statistical analysis. **b** Comparison between nucleotides added during transcription initiation from "AAA" or "TTT" trinucleotides TIRs, as determined by 5'TNET-seq and in vitro transcription. The 5'TNET-seq and in vitro transcription (200 μM NTP) results are indicated by black and blue line, respectively. **c** Boxplot illustrating the mean nucleotides added as determined by the in vitro transcription assay. Promoter tested include the *gcvT*, *deoC* and *fdoG* promoters (AAA, n = 3), and *zapA*, *ppiD* and *xanP* promoters (TTT, n = 3). The cross denotes the mean. Statistical analysis was conducted using a two-tailed student *t*-test.

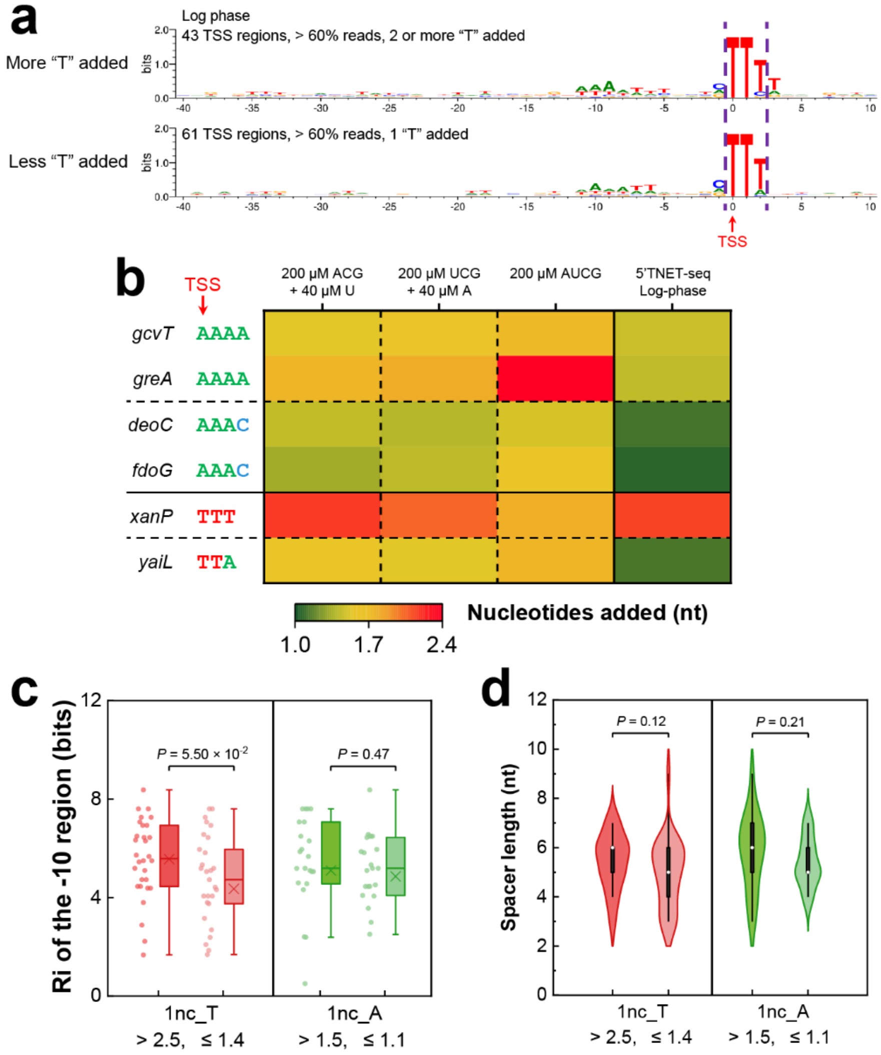


**Supplementary Fig. 12 | Influence of promoter elements on nucleotide addition via productive reiterative initiation. a** Sequence logos of TSS regions with varied numbers of "T" nucleotide added. The first three nucleotides from the TSS are demarcated by dashed lines. **b** Heatmap displaying the number of nucleotides added within transcription initiation regions with varying lengths of homopolymeric tracts. The average number of added nucleotides was calculated based on sequencing data obtained from in vitro transcription assays. Boxplot and violin plot illustrating the information content of the –10 region (**c**) and spacer length (**d**) for 1nc promoters with varying numbers of "T" or "A" nucleotides added. 1nc_T, > 2.5 "T" nucleotides added (n = 30); 1nc_T, ≤ 1.4 "T" nucleotide added (n = 29); 1nc_A, > 1.5 "A" nucleotide added (n = 23); 1nc_A, ≤ 1.1 "A" nucleotide added (n = 23). Statistical analysis was performed using the two-tailed Mann-Whitney *U*-test.

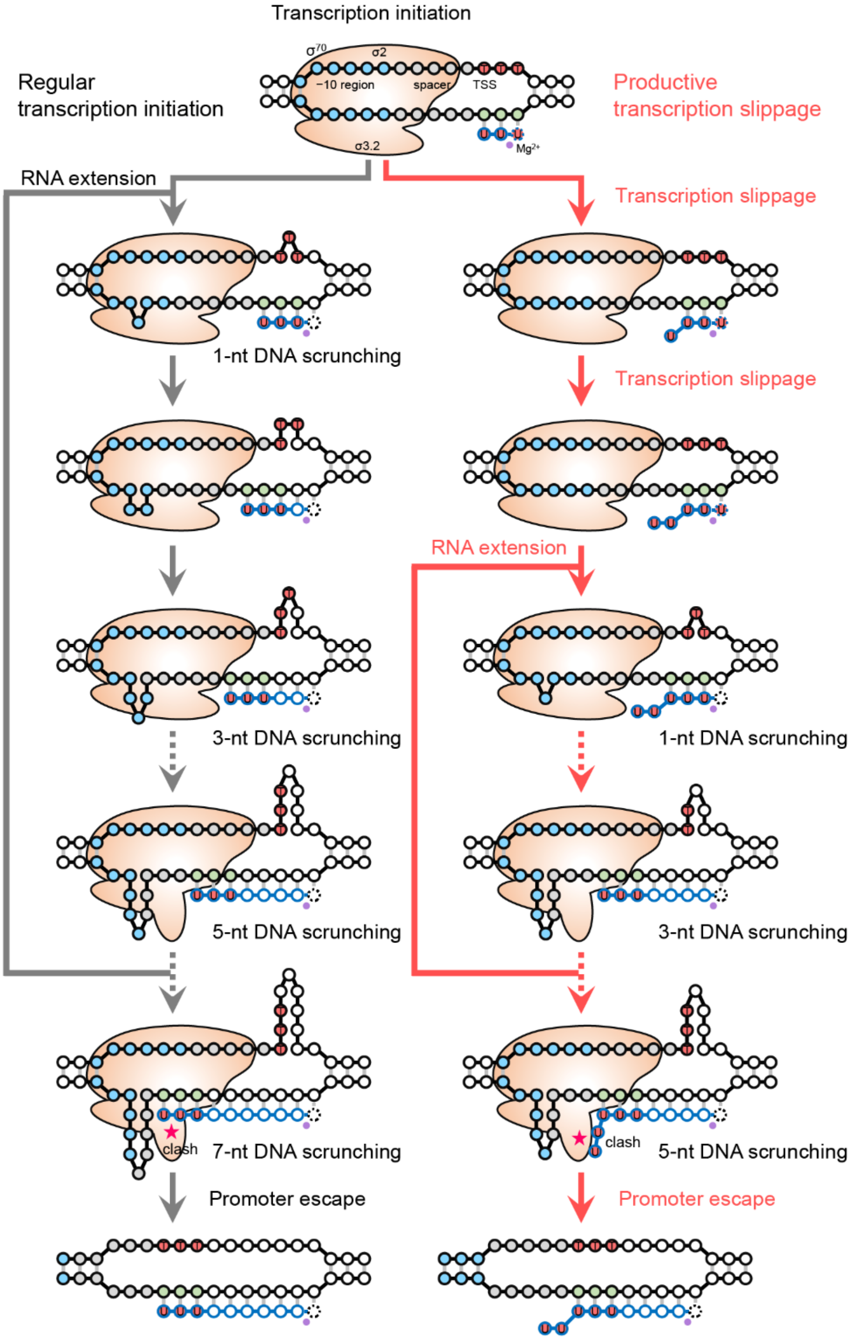


**Supplementary Fig. 13 | Mechanism of enhancing gene transcription through productive reiterative initiation.** The contrasting processes of regular transcription initiation (left) and productive reiterative initiation (right) are indicated by grey and red arrows, respectively. Blue circles, −10 region; grey circles, spacer; red circles, transcription initiation region with trinucleotide "TTT". The bulged nucleotides within the unwound transcription bubble indicate scrunching. Additionally, the clash between the nascent RNA 5' end and σR3.2 is indicated by an asterisk. Only the pretranslocated state of the transcription complexes are shown.

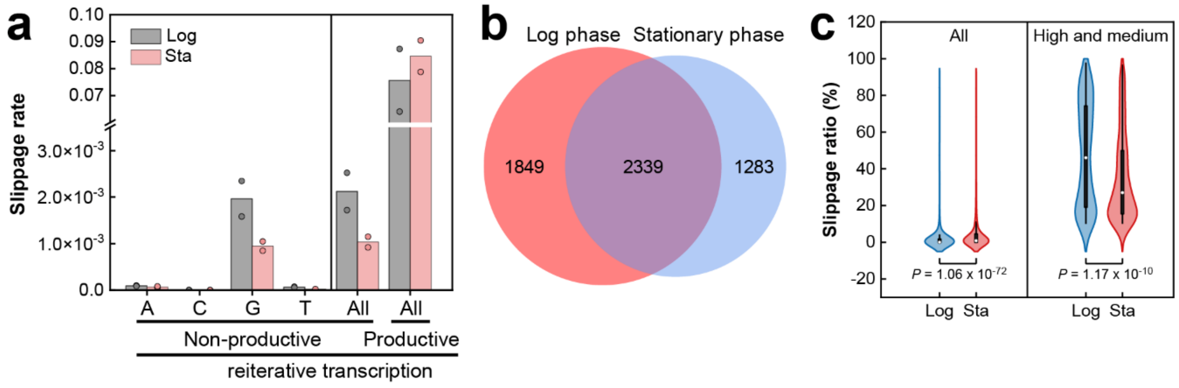


**Supplementary Fig. 14 | Comparative analysis of productive reiterative initiation between log phase and stationary phase. a** Comparison of slippage rates between reads derived from non-productive and productive reiterative initiation, as calculated from 5'TNET-seq data. Sequencing reads that ≥ 80% of the first 15 nucleotides are the same are treated as reads generated by non-productive reiterative initiation. **b** Venn diagram illustrating the intersection of TSSs analyzed at log phase and stationary phase. **c** Violin plot illustrating the slippage ratio for all promoters, as well as those with high and medium slippage at log phase and stationary phase. All-Log, n = 4188; All-Sta, n = 3622; High and medium-Log, n = 425; High and medium-Sta, n = 581. Statistical analysis was performed using a two-tailed Mann-Whitney *U*-test.

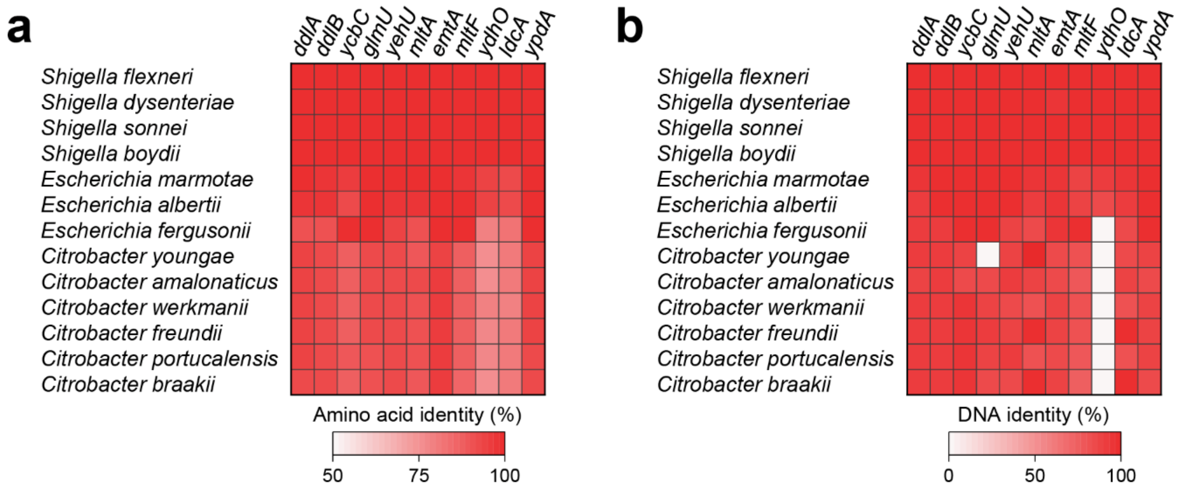


**Supplementary Fig. 15 | Conservation analysis of genes related to cell wall synthesis regulated by productive reiterative initiation in bacteria. a** Heatmap illustrating the conservation of the protein sequences of 11 genes associated with cell wall synthesis across various bacteria, excluding *E. coli*. The entire protein sequences of each gene were subjected to blast analysis to identify the 1000 best-aligned sequences. Subsequently, 13 bacteria containing homologs of all 11 genes were selected for further analysis. **b** Heatmap demonstrating the conservation of regions downstream of the TSSs of the 11 genes. The first 100-bp DNA sequences right downstream of the TSSs were used for blast analysis in the selected 13 bacteria.
